# Reversal of Biological Age in Multiple Rat Organs by Young Porcine Plasma Fraction

**DOI:** 10.1101/2023.08.06.552148

**Authors:** Steve Horvath, Kavita Singh, Ken Raj, Shraddha Khairnar, Akshay Sanghavi, Agnivesh Shrivastava, Joseph A. Zoller, Caesar Z. Li, Claudia B. Herenu, Martina Canatelli-Mallat, Marianne Lehmann, Siniša Habazin, Mislav Novokmet, Frano Vučković, Leah C. Solberg Woods, Angel Garcia Martinez, Tengfei Wang, Priscila Chiavellini, Andrew J. Levine, Hao Chen, Robert T Brooke, Juozas Gordevicius, Gordan Lauc, Rodolfo G. Goya, Harold L. Katcher

**Affiliations:** Department of Human Genetics, David Geffen School of Medicine, University of California, Los Angeles, Los Angeles, California, USA; Department of Biostatistics, Fielding School of Public Health, University of California, Los Angeles, Los Angeles, California, USA; Altos Labs, Cambridge, UK; Shobhaben Pratapbhai Patel School of Pharmacy and Technology Management, SVKM’S NMIMS University, Mumbai, India; Yuvan Research Inc, Mountain View, CA, USA; Institute for Experimental Pharmacology of Cordoba (IFEC), School of Chemical Sciences, National University of Cordoba, Cordoba, Argentina; Biochemistry Research Institute of La Plata – Histology B, Pathology B, School of Medicine, University of La Plata, La Plata CC 455 (zip 1900), Argentina; Genos Glycoscience Research Laboratory, Zagreb, Croatia; Wake Forest University School of Medicine, 1 Medical Center Drive, Winston Salem, NC 27157, USA; Department of Pharmacology, Addiction Science and Toxicology, The University of Tennessee Health Science Center, Memphis, TN 3993, USA; Department of Neurology, David Geffen School of Medicine at the University of California, Los Angeles, CA, 90095, USA; Epigenetic Clock Development Foundation, Torrance, California, USA; Faculty of Pharmacy and Biochemistry, University of Zagreb, Zagreb, Croatia

**Author notes:** Joint first authorship (Steve Horvath, Kavita Singh, Ken Raj). Correspondence: Steve Horvath Harold Katcher.

**Keywords:** rejuvenation, plasma fraction, epigenetic clock, DNA methylation, glycans, rat

## Abstract

Young blood plasma is known to confer beneficial effects on various organs in mice and rats. However, it was not known whether plasma from young pigs rejuvenates old rat tissues at the epigenetic level; whether it alters the epigenetic clock, which is a highly accurate molecular biomarker of aging. To address this question, we developed and validated six different epigenetic clocks for rat tissues that are based on DNA methylation values derived from n=613 tissue samples. As indicated by their respective names, the rat pan-tissue clock can be applied to DNA methylation profiles from all rat tissues, while the rat brain-, liver-, and blood clocks apply to the corresponding tissue types. We also developed two epigenetic clocks that apply to both human and rat tissues by adding n=1366 human tissue samples to the training data. We employed these six rat clocks to investigate the rejuvenation effects of a porcine plasma fraction treatment in different rat tissues. The treatment more than halved the epigenetic ages of blood, heart, and liver tissue. A less pronounced, but statistically significant, rejuvenation effect could be observed in the hypothalamus. The treatment was accompanied by progressive improvement in the function of these organs as ascertained through numerous biochemical/physiological biomarkers and behavioral responses to assess cognitive functions. An immunoglobulin G (IgG) N-glycosylation pattern shift from pro-to anti-inflammatory also indicated reversal of glycan aging. Overall, this study demonstrates that a young porcine plasma-derived treatment markedly reverses aging in rats according to epigenetic clocks, IgG glycans, and other biomarkers of aging.

## INTRODUCTION

The field of heterochronic parabiosis, involving the mixing of blood from young and old mice, laid the groundwork for investigations into the impacts of exposing an aged organism to a youthful systemic environment. Originating in rats before expanding to mice, this field was initially pioneered by Clive McCay who employed it in the study of aging in rats, although early attempts were hampered by the widespread occurrence of ’parabiotic disease’ ^1, 2^. Progressing through time, surgical methods improved, and rat studies by Ludwig and Elashoff (1972) unveiled the lifespan extension of an older parabiont when partnered with a younger one, presenting the first evidence of extended life in the older organism as a response to a youthful environment ^3^.

Later investigations using mice revealed insights into aging physiology and stem cell behavior across different tissues and organ systems ^4, 5^. Beneficial influences of young blood on various murine organs were evident in functional tests, with outcomes suggesting that the beneficial effects were a result of blood-borne factors as opposed to indirect benefits from the young mice’s superior organ functionality ^6–12^. The ensuing interest led to widespread research and commercial attention on the identification and isolation of rejuvenation factors from blood that could potentially mitigate or treat age-related conditions.

However, in the context of aging and rejuvenation, it is crucial to differentiate between improved health or organ function, which could be achieved via medication or surgery, and genuine molecular age reversal. Clinical biomarkers, despite their utility in indicating organ dysfunction or disease, fall short of accurately indicating fundamental aging mechanisms ^13^. With this in mind, our study tackles the question of whether plasma fraction treatment or similar interventions can truly reverse biological age, using two molecular aging biomarkers – DNA methylation and glycation.

Epigenetic changes, including cytosine methylation, are a well-established hallmark of aging, useful for developing accurate age estimators ^14–21^. Notably, DNA methylation (DNAm) age estimators apply broadly, including to sorted cells, tissues, and organs, across the entire age spectrum. The difference between DNAm age and chronological age, termed “epigenetic age acceleration”, is predictive of all-cause mortality and is associated with a wide variety of conditions ^22–25, 26^^27–30, 2015^.

Complementing the epigenetic clocks, analyses of human immunoglobulin G (IgG) have established that changes in its N-glycome composition occur with aging and disease, providing a ’glycan clock’ that can indicate age and be reversed through lifestyle changes ^31, 32^. Epigenetic clocks have been successfully used to evaluate age-related interventions in mice, and recently, we developed six epigenetic clocks for rats, two of which are applicable to humans as well ^33–37^.

This paper aims to address the question: will young plasma from pigs reverse the epigenetic age of rat tissues? To assess this, we used these six rat-specific epigenetic clocks along with IgG N-glycan profiling, physiological, histological, biochemical and cognitive assessments, to evaluate the plasma fraction-based treatment in tissues from 2-year-old rats. Our findings demonstrate that this approach greatly reverses aging in rats.

## RESULTS

### DNA methylation data

All DNA methylation data were generated on a custom methylation array that applies to all mammals. We obtained in total, DNA methylation profiles of n=613 samples from 13 different tissues of rat (*Rattus norvegicus*) (**Supplementary Table S1, Supplementary Table S2, Supplementary Table S3**), with ages that ranged from 0.0384 years (i.e., 2 weeks) to 2.3 years (i.e., 120 weeks). The rat tissue samples were from 3 different countries: (i) India (Yuvan Research in collaboration with NMIMS School of Pharmacy’s), (ii) the United States (H. Chen and L. Solberg Woods), and (iii) Argentina (R. Goya). Unsupervised hierarchical clustering shows that the methylation profiles clustered by tissue type, as would be expected (**Supplementary Figure 1**).

Our DNA methylation-based age estimators (epigenetic clocks) were developed (“trained” in the parlance of machine learning) using n=503 rat tissues. The two epigenetic clocks that apply to both species were developed by adding n=1366 human tissue samples to the rat training set. Both rat and human tissues were profiled on the same methylation array platform (HorvathMammalMethylChip40) that focuses on 36,000 highly conserved CpGs ^38^ (**Methods**).

### Epigenetic clocks

Our six different clocks for rats can be distinguished along several dimensions (tissue type, species, and measure of age). Some clocks apply to all tissues (pan-tissue clocks) while others are tailor-made for specific tissues/organs (brain, blood, liver). The rat pan-tissue clock was trained on all available tissues. The brain clock was trained using DNA samples extracted from whole brain, hippocampus, hypothalamus, neocortex, substantia nigra, cerebellum, and the pituitary gland. The liver and blood clock were trained using the liver and blood samples from the training set, respectively. While the four rat clocks (pan-tissue-, brain-, blood-, and liver clocks) apply only to rats, the human-rat clocks apply to both species. The two human-rat pan-tissue clocks are distinct, by way of measurement parameters. One estimates *absolute* age (in units of years), while the other estimates *relative* age, which is the ratio of chronological age to maximum lifespan; with values between 0 and 1. This ratio allows alignment and biologically meaningful comparison between species with very different lifespan (rat and human), which is not afforded by mere measurement of absolute age.

To arrive at unbiased estimates of the six epigenetic clocks, we used a) cross-validation of the training data and b) evaluation with an independent test data set. The cross-validation study reports unbiased estimates of the age correlation R (defined as Pearson correlation between the age estimate (DNAm age) and chronological age) as well as the median absolute error (**Figure 1**). The cross-validation estimates of the age correlations for all six clocks are higher than 0.85. The four rat clocks exhibited median absolute errors that range from 0.137 years (1.6 months) for the rat blood clock to 0.182 years (2.2 months) for the rat pan-tissue clock, **Figure 1A-D**). The human-rat clock for age generated an age correlation of R=0.99 when both species are analyzed together (**Figure 1E**) but is lower when the analysis is restricted to rat tissues alone (R=0.85, **Figure 1F**). In contrast, the human-rat clock for *relative age* exhibits high correlation regardless of whether the analysis is done with samples from both species (R=0.96, **Figure 1G**) or only with rat samples (R=0.94, **Figure 1H**). This demonstrates that relative age circumvents the skewing that is inherent when absolute age of species with very different lifespans are measured using a single formula. This is due in part to the unequal distribution of training data at the opposite ends of the age range.

**Figure 1:**
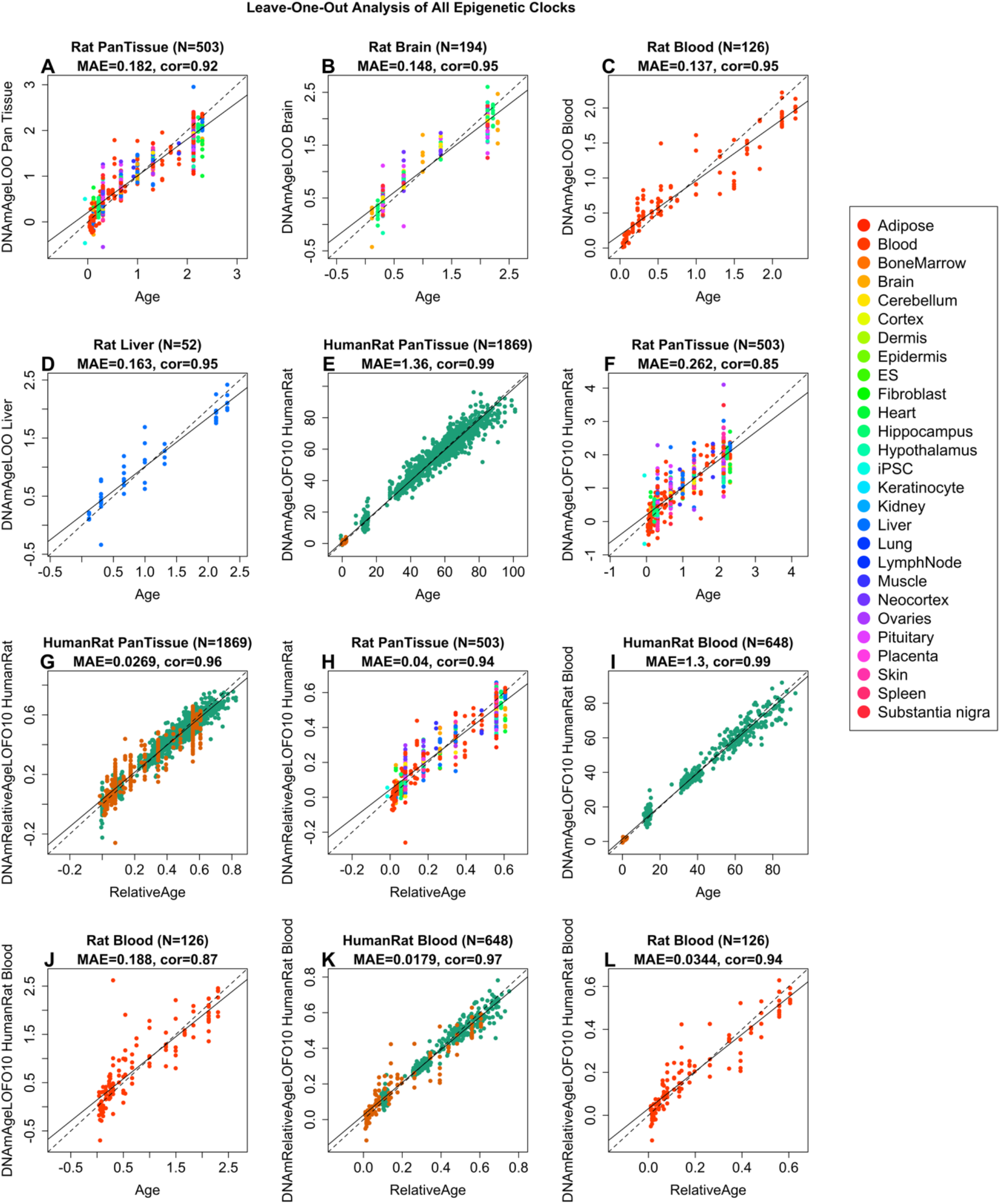
Cross-validation study of six epigenetic clocks for rat. A-D) Four epigenetic clocks that were trained on rat tissues only. E-H) Results for 2 clocks that were trained on both human and rat tissues. Leave-one-sample-out estimate of DNA methylation age (y-axis, in units of years) versus chronological age for A) Rat pan-tissue, B) Rat brain, C) Rat blood, and D) Rat liver clock. Dots are colored by A) tissue type or B) brain region. E and F) “Human-rat” clock estimate of absolute age. G, H) Human-rat clock estimate of relative age, which is the ratio of chronological age to the maximum lifespan of the respective species. Ten-fold cross-validation estimates of age (y-axis, in years) in E, G) Human (green) and rat (orange) samples and F, H) rat samples only (colored by tissue type). Each panel reports the sample size, correlation coefficient, median absolute error (MAE).

As indicated by its name, the rat pan-tissue clock is highly accurate in age estimation of all the different tissue samples tested and its performance with individual tissues can be more clearly seen in **Supplementary Figure 2**. We also evaluated the accuracy of the six epigenetic clocks in independent test data from the plasma fraction test study. In the untreated rat tissue samples, the epigenetic clocks exhibited high age correlations in all tissues (R>=0.95 in blood, liver, and the hypothalamus and R>=0.89 in Heart Tissue, **Supplementary Figure 3**).

### Rejuvenation effect of the plasma fraction treatment

The ability to generate epigenetic age clocks for specific rat organs can be readily appreciated from the perspective of their practical utility in aging research. However, the appreciation of a pan-tissue clock is much deeper as it extends into the conceptual aspect with regards to aging. The ability to estimate the age of different rat organs with a single clock; be it the rat pan-tissue clock or the human-rat pan-tissue clock, is a very strong indicator that epigenetic age is regulated across all tissues of an organism, and this regulation is mediated systemically. This in turn implies that it may be possible to alter the rate of aging centrally and simultaneously across different tissues of the body; a principle that underlies the plasma fraction treatment.

The plasma fraction treatment used in the investigation described below, is based on the principle of Heterochronic Plasma Exchange (HPE) ^39^, which does not involve the physical attachment of the circulatory systems of two animals. In addition to greatly reducing the stress to the animals, this is expected to have a more profound effect as 100% of the old animal’s blood could in theory be replaced. This contrasts with heterochronic parabiosis, where a young rat, with approximately half the weight of an old rat, contributes less than 50% of the combined plasma circulation in the parabiotic partners. The general loss of tissue repair with age may reflect the negative influence of age-accumulated inhibitory proteins in aged tissues and circulation ^40^. Therefore, if putative pro-aging factors are present in the plasma of old animals ^40^, they would be present in the old parabiotic partner and be distributed to the young one, causing it to age more rapidly.

The plasma fraction treatment described here is a step change from HPE, as it uses neither whole blood nor plasma, but the exosome fraction of the plasma, which we term as E5 (**Methods**). Another significant novelty is the source of E5, which were piglets. The significance and implications of this are elaborated in the discussion section below. We applied the six clocks to an independent test data set (n=76) comprising four rat tissues (blood, liver, heart, hypothalamus) to test whether a porcine plasma fraction from 6-month-old piglets, injected into 2-year-old rats would reverse their epigenetic age.

Plasma fraction treatment was administered to male rats following the experimental plan depicted in **Supplementary Figure 4.** Briefly, 18 male Sprague Dawley rats were divided into three groups. A group of 6 young rats (30 weeks old), a second group of 6 old rats (109 weeks old) and a third group of 6 plasma fraction-treated old rats (also 109 weeks old). Plasma fraction treatment consists of two series of intravenous injections of E5. Rats were injected four times on alternate days for 8 days. A second identical series of injections were administered 95 days later. In its entirety, the experiment lasted 155 days. For the duration of the experiment, blood was drawn at regular intervals for hematological and biochemical analyses to monitor the impact of the treatment on blood, and solid vital organs. Cognitive functions of the rats were assessed four times during this period and at the end of the experiment, the animals were sacrificed, and DNA methylation profiles of several organs were generated. The DNA methylation profiles were evaluated using the above-mentioned six epigenetic clocks. The results derived from these profiles are plotted in **Figure 2**, which shows that the epigenetic ages of the old and young rats are readily distinguishable by all the six clocks. Crucially, plasma treatment of the old rats reduced the epigenetic ages of blood, liver and heart by a very large and significant margin, to levels that are comparable with those of the young rats. According to the 6 epigenetic clocks, the plasma fraction treatment rejuvenated liver by 74.6% (ranging from 68.6% to 78.6% depending on the clock, Supplementary Table S4), blood by 64.3% (ranging from 52.5 to 74.5%), heart by 46.5%, and hypothalamus by 24.4%. It is necessary to point out that the epigenetic clocks that were employed to analyze these experimental samples were developed independently of any of the methylation data from the experimental samples. This is important as it establishes that the E5-induced rejuvenation that were observed, were not influenced by the way these clocks were developed. After having convincingly demonstrated that, we included DNA methylation profiles from the untreated controls of the experiment to the training set, to derive an even more accurate clock (elaborated in the section below). Epigenetic age measurements of the experimental samples with the resulting epigenetic clocks show rejuvenation effects of E5 that are even more pronounced: liver 77.6%, blood 68.2%, heart 56.5%, hypothalamus 29.6%, with the average rejuvenation across four tissues of 67.40%. In other words, the treatment more than halved the epigenetic age (**Figure 2I-P**).

**Figure 2:**
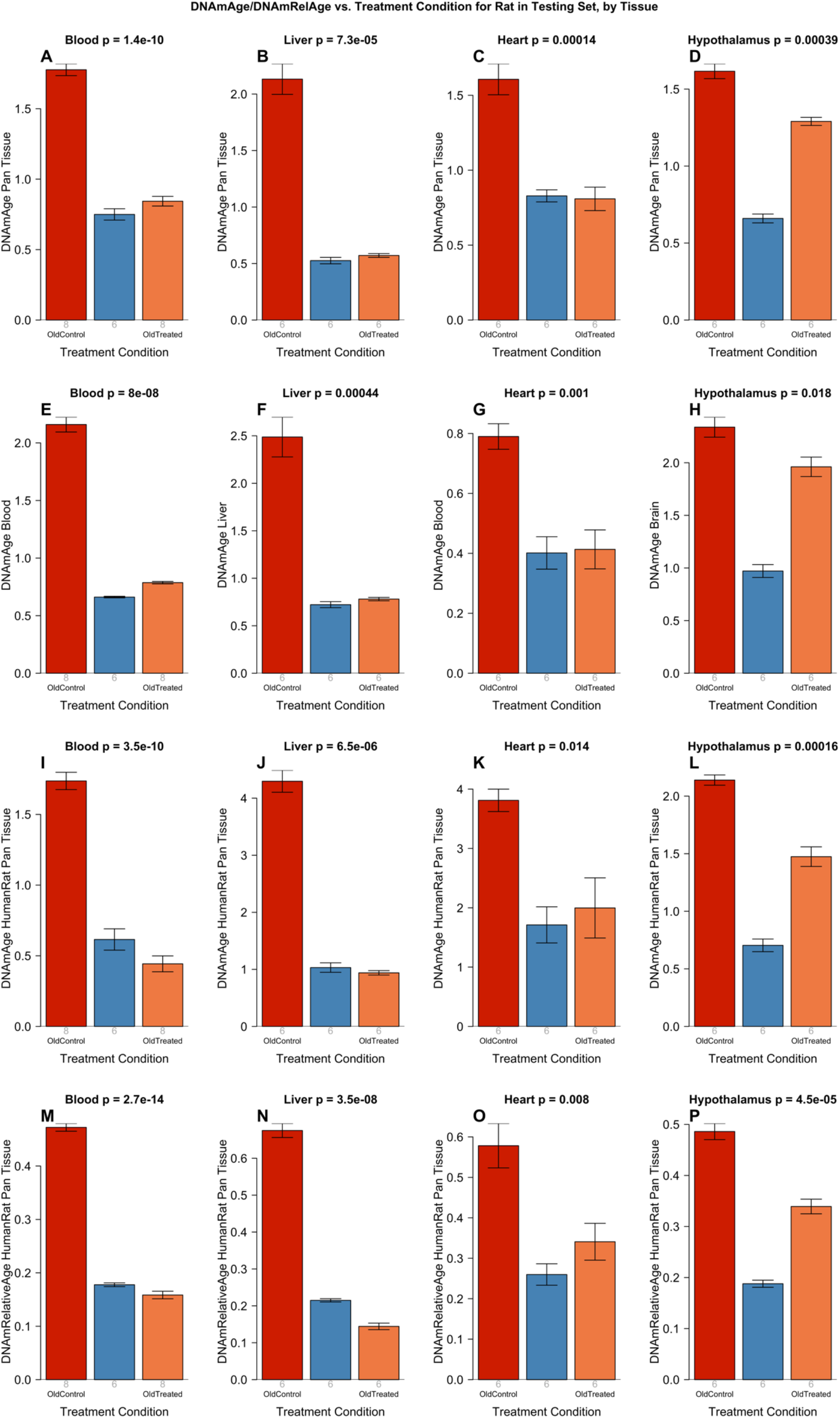
Epigenetic clock analysis of plasma fraction treatment. Six epigenetic clocks applied to independent test data from four rat tissue type (columns): blood, liver, heart, and hypothalamus. A-D) Rat pan-tissue clock. E) Rat blood clock applied to blood. F) Rat liver clock applied to liver. G) Rat blood clock applied to heart. H) Rat brain clock applied to hypothalamus. I-L) Human-rat clock measure of absolute age. M-P) Human-rat clock measure of relative age defined as age/maximum species lifespan. Each bar-plot reports the mean value and one standard error. P values results from analysis of variance. Student T-test p values result from a 2-group comparison of old controls (left bar) versus old treated samples (right bar), i.e. the young controls were omitted.

### Final version of epigenetic clocks

These final versions of clocks were developed by adding the “untreated” samples from the E5 test data to the original training data (n=503 rat tissues). This increased sample size of the training data, led to clocks with higher accuracy, as shown by a cross validation analysis (**Supplementary Figure 5, Supplementary Figure 6**). Using this final version of the epigenetic clocks, we find that the treatment effects were even more significant especially for the hypothalamus **(Supplementary Figure 7**). Final versions of the pan-tissue clock, liver clock, blood clock, brain clock, and “human-rat” clock can be found in Supplementary Material. These clocks would be of great use in future experiments with rats.

### EWAS of age and treatment effects

Epigenetic clocks are attractive aggregate biomarkers because they summarize the information of many CpGs into a single number (the age estimate). They are nevertheless products of only a subset of age-related CpG changes. Hence, it is also informative to look at all CpGs in response to treatment. We executed an epigenome-wide association study, employing a false discovery rate threshold of 0.05 for both aging and treatment effects (refer to Methods for details). A strong reversal in age-related methylation gain post-E5 treatment was observed in blood, liver, heart, and the hypothalamus, as presented in **Supplementary Figure 8**. In summary, E5 treatment effectively reversed the alterations in methylation at individual CpGs, which typically occur with aging.

#### Physical and overt effects

The reduction of epigenetic age of E5-treated rats is particularly significant as it would appear to indicate that aging is a coordinated process as opposed to a stochastic one that occurs independently between the different organs. Before any further consideration of this notion, it is necessary to determine whether the reduction in epigenetic age is indeed biologically meaningful. In other words, is the rejuvenation of epigenetic age accompanied by changes in other well-characterized age-related endpoints. Equally important is the need to determine whether E5 treatment generated any adverse side-effects.

The weight of rats for the duration of the experiment was monitored at regular intervals and the plots in **Supplementary Figure 9A** indicate that E5 treatment did not affect food intake and appetite in any way, as the weight of the treated and untreated rats were similar, and there was no presentation of any overt signs of physical or behavioral abnormality. These features replicated those we observed in a mid-term (30-day) pilot experiment, where in addition, we also measured and found grip strength of old rats to be considerably improved by this treatment (**Supplementary Figure 9B, Supplementary Table S5**). At 15 days post-treatment, the strength of plasma fraction-treated old rats was indistinguishable from that of young ones. These and other encouraging results from the mid-term pilot study, prompted a longer-term (155-day) investigation, with a new preparation of E5. The result from this study forms the main corpus of this report. These two independent investigations produced similar results that varied only in terms of magnitude, as is consistent with their different durations. Histological examinations of the various organs did not indicate any obvious abnormalities after 155 days of treatment (**Supplementary Figure 10** and **Supplementary Table S6**). Instead, oil red O staining showed that accumulation of fat in old tissues was greatly reduced in E5-treated rats (**Supplementary Figure 11**).

### Blood cell indices and hematology

To monitor potential effects of E5 on the blood of rats, we measured hemoglobin levels, mean corpuscular volume (MCV), mean corpuscular hemoglobin (MCH), mean corpuscular hemoglobin concentration (MCHC), hematocrit (HCT) levels and obtained counts of red blood cells, platelets, white blood cells and lymphocytes at 0, 60 and 155 days from the start of treatment. These blood indices are informative indicators of malfunction of bone marrow and vital organs and importantly, they vary with the age of the animal. In, all these blood indices were very different between young and old rats at the start of the experiment, and in time, plasma fraction treatment nudged all these parameters, except for platelets, away from the values exhibited by old untreated rats towards those of the young ones (**Supplementary Figure 12** and **Supplementary Table S7**). This is easily visualized by the movement of the orange dots, representing treated old rats in each graph towards the blue dots, representing young rats. Plasma fraction treatment has not caused any changes to the blood indices that would indicate any organ dysfunction. Instead, it rejuvenated the blood of the rats, which is consistent with the significant reduction of epigenetic age of their blood as measured by both the rat multi-tissue and blood clocks.

### Biomarkers for vital organs

To ascertain the impact of plasma fraction treatment on vital organs, we measured the levels of the following biomarkers on 30, 60, 90, 120 and 155 days from the start of the experiment: bilirubin, serum glutamic-pyruvic transaminase (SGPT) and serum glutamic-oxaloacetic transaminase (SGOT) to monitor liver function; triglycerides (TG), HLD and cholesterol to monitor risk of atherosclerosis and heart disease, and liver function as well; glucose to monitor the pancreas and diabetes; and creatinine and blood urea nitrogen for kidney function. The levels of all these biomarkers in the treated old rats were altered towards the values of young rats, without exception **(Figure 3** and **Supplementary Table S8)**. This is easily visualized by the movement of the orange dots, representing treated old rats in each graph towards the blue dots, which represent young rats. Collectively, these results show that the function of all the vital organs tested through their respective biomarkers, were rejuvenated by E5 treatment. This is entirely consistent with reversal of the epigenetic ages of their hearts and livers.

**Figure 3:**
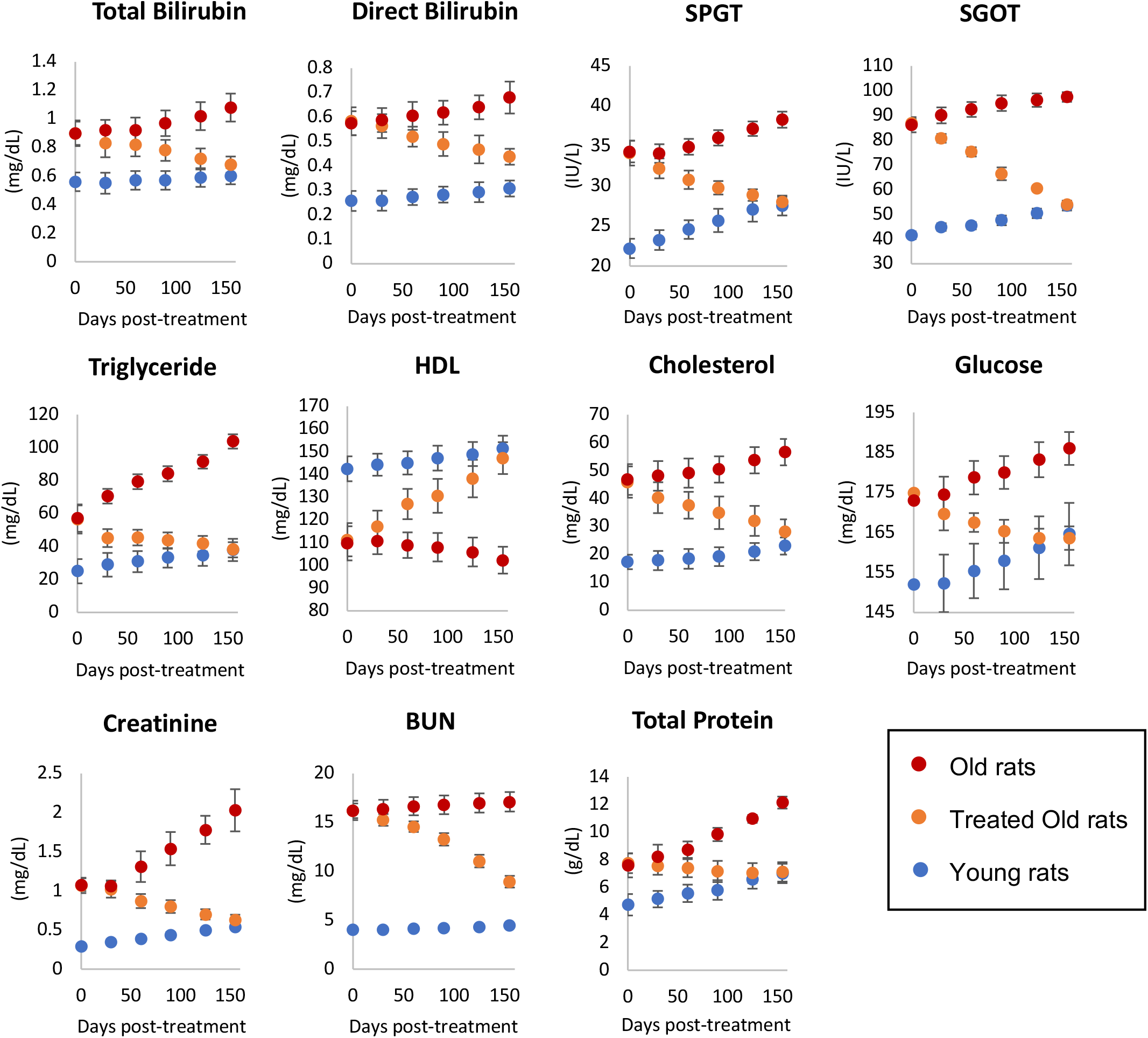
Assessment of plasma fraction treatment on vital organ functions. At 0, 30, 60, 90, 120 and 155 days from commencement of the experiment, the health and function of vital organs (liver, heart, kidney and pancreas) of 6 rats per group were monitored through measurements of appropriate biomarkers. SPGT = serum glutamic-pyruvic transaminase, SGOT = serum glutamic-oxaloacetic transaminase, HDL = high-density lipoprotein and BUN = blood urea nitrogen. Red dots represent data points of old rats, orange dots represent treated old rats and blue represent young rats. Each data point represents average values from 6 rats with 2 standard errors around the mean. Detailed measurements of each parameter are provided in Supplementary Table S8.

### Cognitive function

Learning and memory, which are constituent characteristics of cognitive functions, decline not only in humans but also in rats, starting from 12 months of age ^38^. Barnes maze was used to measure the latency period required by the rats to escape through the right hole into an escape box. Videos illustrating the latency pattern of three rats (young, old treated, old untreated) can be found in the Supplement. Within a month of plasma fraction treatment, the rats exhibited significantly reduced latency to escape (**Figure 4**), i.e., they learned and remembered better. After the second month, the treated rats began with a slightly reduced latency period compared to the untreated old rats, and once again, they learned much faster than the latter. By the third month, it was clear that treated rats remembered the maze much better than the untreated ones even from the first day of test as their latency period was significantly reduced and by the end of the test period their latency was like that of the young rats. This feature was sustained and repeated in the fourth month. Collectively, these results show that E5 improved the learning and memory of the rats. Interestingly, the epigenetic age of treated rat brain samples (hypothalamus) was lower than the untreated old ones, but less markedly than the magnitude of decrease of epigenetic age of the blood, heart and liver. This introduces numerous possible insights into the relationship between cognitive function, biological age and physical health which will be further elaborated in the discussion.

**Figure 4:**
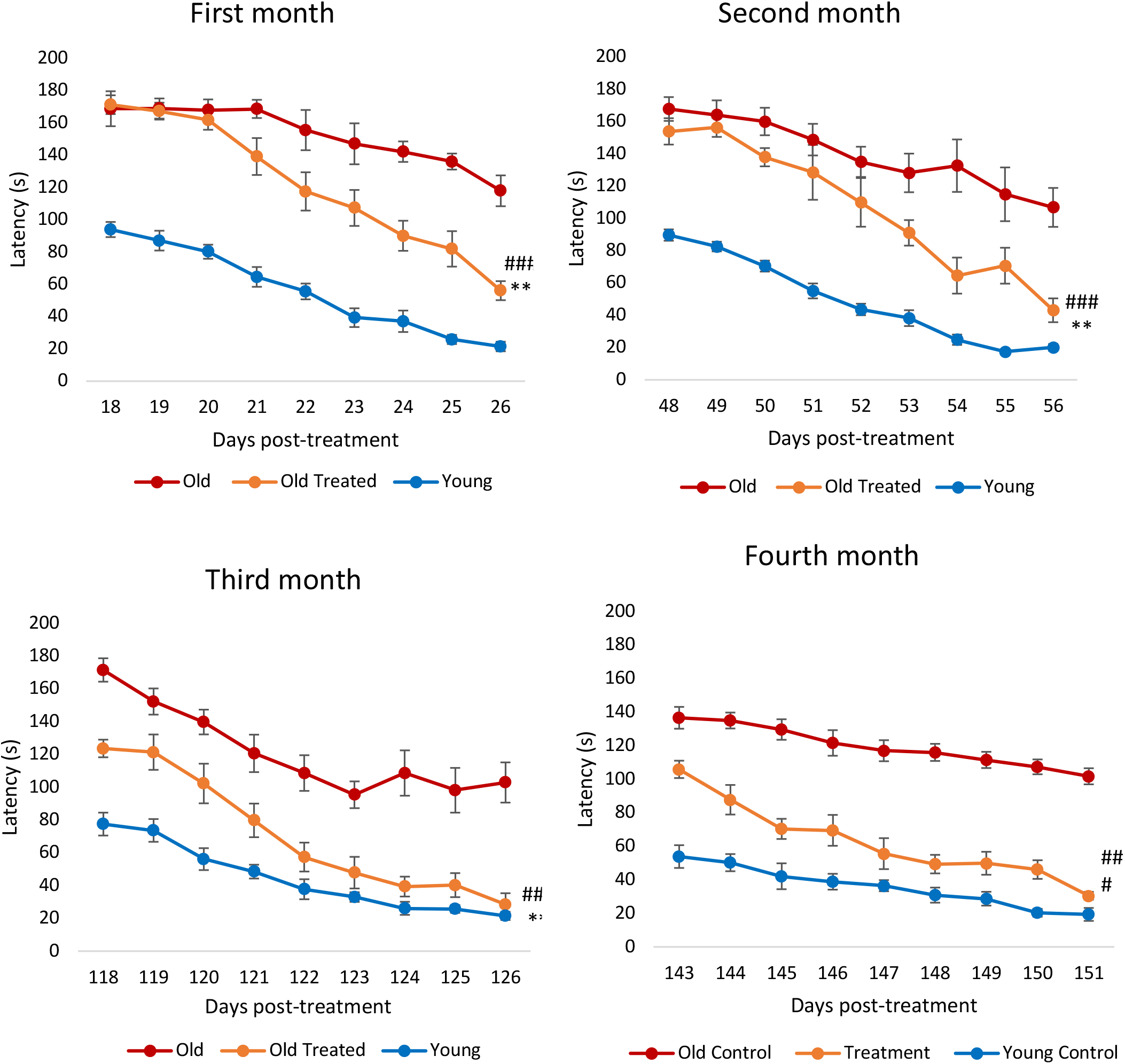
Assessment of plasma fraction treatment on cognitive function (learning and memory). Rats were subjected to Barnes maze test in the first to fourth month from commencement of the experiment. Each assessment consists of nine consecutive days of test where the time (in seconds) required by the rats to find the escape hole (latency) was recorded and plotted. The error bars depict 2 standard errors.

### Cellular stress

There is no doubt that the reduction in epigenetic age of liver, heart, brain and blood by plasma fraction was accompanied by startling improvement in the function of these organs. In addition to the decline of organ function with age, is the rise of two cell stress features which are related, namely oxidative stress and chronic inflammation; the excess of which have been linked to multiple pathologies. We carried out a panel of biomarker tests for these two features in the test rats.

#### Oxidative stress

Oxidative stress results from the excess of reactive oxygen species (ROS) in cells. This situation can arise due to the over-production of ROS or the decline in the ability to remove or neutralize ROS. While ROS at low levels is not harmful and are even necessary, at higher levels, they interact with biomolecules such as lipids and compromise their function. Measuring the levels of malondialdehyde (MDA), which is the end-product of poly-unsaturated fatty acid peroxidation, reveals the levels of cellular ROS. The amounts of MDA were clearly higher in the brain, heart, lung and liver of older rats (**Figure 5**), and plasma fraction treatment reduced this level to that of young rats. Hence, regardless of the source of augmented ROS in older rats, plasma fraction appears to be able to reduce it effectively.

**Figure 5:**
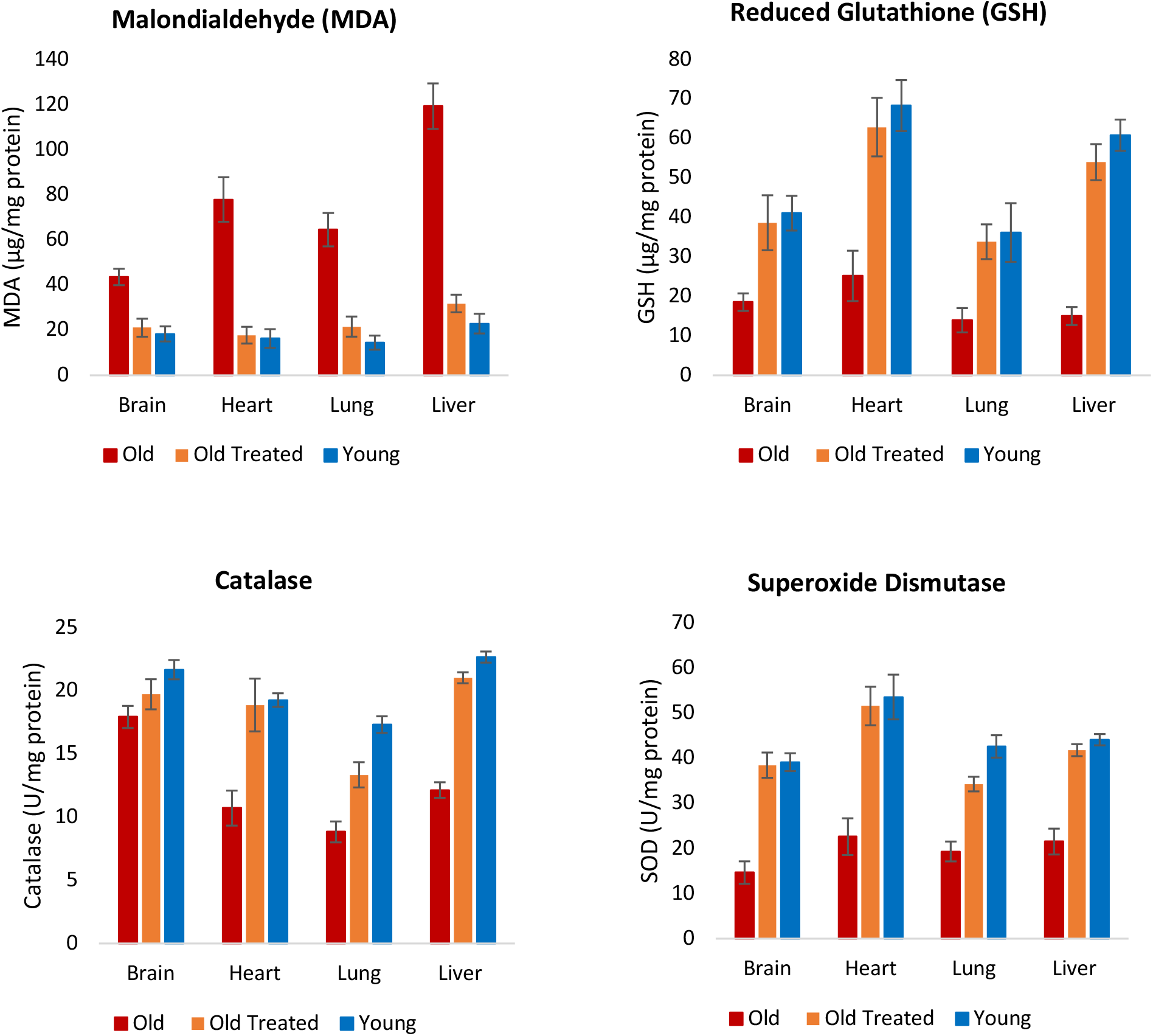
Assessment of plasma fraction treatment on oxidative stress. At the end of the experimental period of 155 days, lipid peroxidation level, which is an indicator of intracellular level of reactive oxygen species (ROS) was determined by measuring the amount of Malondialdehyde (MDA), which is the endproduct of polyunsaturated fatty acid peroxidation. The levels of three antioxidants; reduced glutathione, catalase and superoxide dismutase levels were also measured to ascertain the impact of plasma fraction treatment on oxidative stress. The error bars depict 2 standard errors.

Apart from increased production of ROS, decreased efficiency in eliminating ROS also contributes to the age-associated rise in its level. ROS are neutralized by cellular antioxidants including but not limited to reduced glutathione, catalase and superoxide dismutase, which all work in very different ways. The levels of these three antioxidants were reduced in tissues of old untreated rats but the plasma fraction treatment augmented them to levels that are comparable to young ones (**Figure 5**).

It remains to be ascertained if the reduction of ROS levels, as measured by MDA, was due entirely to the increase in the amounts of antioxidants, or whether plasma fraction treatment also induced a concomitant reduction in the production of ROS from multiple and various intracellular sources. Nevertheless, the endpoint of ROS levels in old rats being diminished to the level of young ones is yet another indication of the rejuvenating effect of the treatment.

#### Chronic inflammation

Inflammation is an important response that helps protect the body, but excess inflammation especially in terms of duration of this response can have very detrimental effects instead. This occurs when inflammation fails to subside and persists indefinitely; a condition referred to as chronic inflammation, which for reasons not well-understood, increases with age and is associated with a multitude of conditions and pathologies. The levels of two of the most reliable and common biomarkers of chronic inflammation; interleukin 6 (IL-6) and tumor necrosis factor α (TNF-α) are found to be considerably higher in old rats (**Figure 6**), and these were very rapidly diminished, within days by E5 treatment, to comparable levels with those of young rats. This was especially stark with IL-6. In time, the levels of these inflammatory factors began to rise gradually, but they were once again very effectively reduced following the second administration of E5 on the 95^th^ day. The reduction of these inflammation markers is consistent with the profile of the nuclear factor erythroid 2-like 2 protein (Nrf2), which plays a major role in resolving inflammation, in part by inhibiting the expression of IL-6 and TNF-α. Nrf2 also induces the expression of antioxidants that neutralizes ROS, which is also a significant feature in inflammation ^41^. In summary, plasma fraction reduces oxidative stress and chronic inflammation, which are age-associated pan-tissue stresses, to the levels found in young rats.

**Figure 6:**
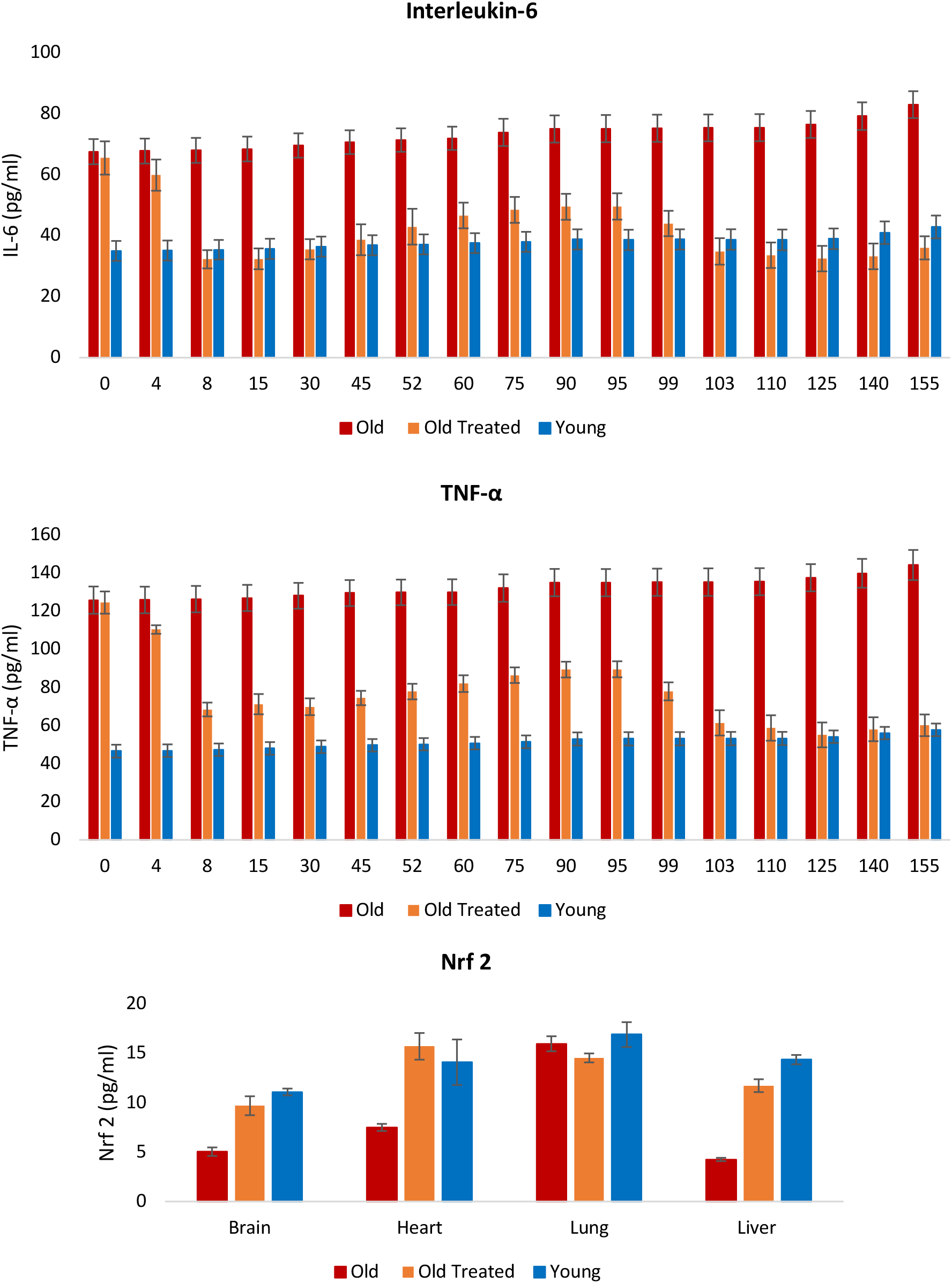
Assessment of the effect of plasma fraction treatment on chronic and systemic inflammation. Blood levels of interleukin-6 (IL-6) and tumor necrosis factor α (TNF-α) were measured at regular intervals throughout the 155-day period of the experiment. At the end of the experiment the levels of Nrf2, a pivotal modulator of inflammation and oxidative stress were measured in brain, heart, lung and liver of the rats. The error bars depict 2 standard errors.

### Additional validation: E5 works in both sexes

The pronounced rejuvenation effects in male rats prompted us to conduct further confirmatory experiments. A particularly important consideration is the effectiveness of E5 with regards to sex, as sex-dependent rejuvenation by some interventions have previously been reported ^42^. To assess E5’s applicability to both male and female Sprague Dawley rats, we studied 12 males (6 treated with E5, 6 with saline) and 12 females (6 treated with E5, 6 with saline). These rats were treated every 45 days with an injection of E5 or saline. The rats were monitored for 165 days, and blood was drawn at six time points: 0, 15, 30, 60, 150 and 165 days from the first injection, (**Supplementary Figure 13**). We observed substantial improvements in IL6 and TNF alpha levels in the blood of both sexes within 15 days of the first injection of E5 (Student T test p<0.05, **Supplementary Figure 13**). At the end of the study (165 days), after 3 dosing cycles of E5, we observed highly significant improvements in TNF alpha levels (Student T test p=1.3×10^-6^ for males and p=7.6×10^-5^ for females) and IL6 levels (p=4.1×10^-6^ for males, p=3.4×10^-6^ for females, Supplementary Figure 13A-D) in the blood of E5-injected rats over that of saline controls. We also observed a substantial improvement in grip strength (p=6.8×10^-7^ for males, p=4.6×10^-6^ for females, **Supplementary Figure 13E,F**). Our study shows age reversal effects in both male and female rats, but E5 is more effective in males.

### Second E5 methylation experiments in blood

To validate our epigenetic clock results, we conducted a second set of E5 experiments with Sprague Dawley rats of both sexes. When these rats turned 26 months old, half (9 rats) received the E5 treatment while the other half (8 rats) received only the control treatment (saline injection). We analyzed methylation data from two blood draws: blood draw before treatment (baseline) and a follow up sample (15 days after the E5/saline treatment).

Again, we observed significant rejuvenation according to the final rat clock for blood (two sided Kruskal Wallis p=0.0094, **Supplementary Figure 14A,B**), and the final pan-tissue clock (p=0.054, Supplementary Figure 14C,D). After omitting an outlying control sample, we obtained more significant results in both sexes (p=0.00086, p=0.014 in females, p=0.053 in males, **Supplementary Figure 14F-J**)

Our results are consistent with the results from a different evaluation of young whole plasma extracted from young rats. In this study, whole plasma from young rats (2 months) was intraperitoneally injected into 25-month-old rats until their natural death ^43^. As in the current study, this separate investigation using whole plasma also found that very soon following treatment, the DNAm age of the treated rats fell below that of controls and remained lower until the end of their lives ^43^.

### Evaluation Immunoglobulin G N-glycan aging

The pivotal function of IgG N-glycosylation in the realms of immunity and inflammation correlates its modifications with inflammaging – a foundational theme in age-related diseases^44^. Until now, the glycan aging clock, informed by the glycomics of IgG, has been employed as an indicator of biological aging exclusively in humans.

In this study, we expanded upon this framework and leveraged glycoproteomics to determine the impact of E5 treatment on the glycan aging of rat IgG. We recorded alterations in rat IgG Fc N-glycosylation for subclasses IgG2a, IgG2b, and IgG2c at two distinct timepoints. Among the control group of untreated rats, we noticed an escalation in IgG agalactosylation (IgG2a-G0) and a reduction in digalactosylation (IgG2a-G2). This overarching pattern mirrors the typical trend observed during human biological aging (**Figure 7** and **Supplementary Table S9**).

**Figure 7:**
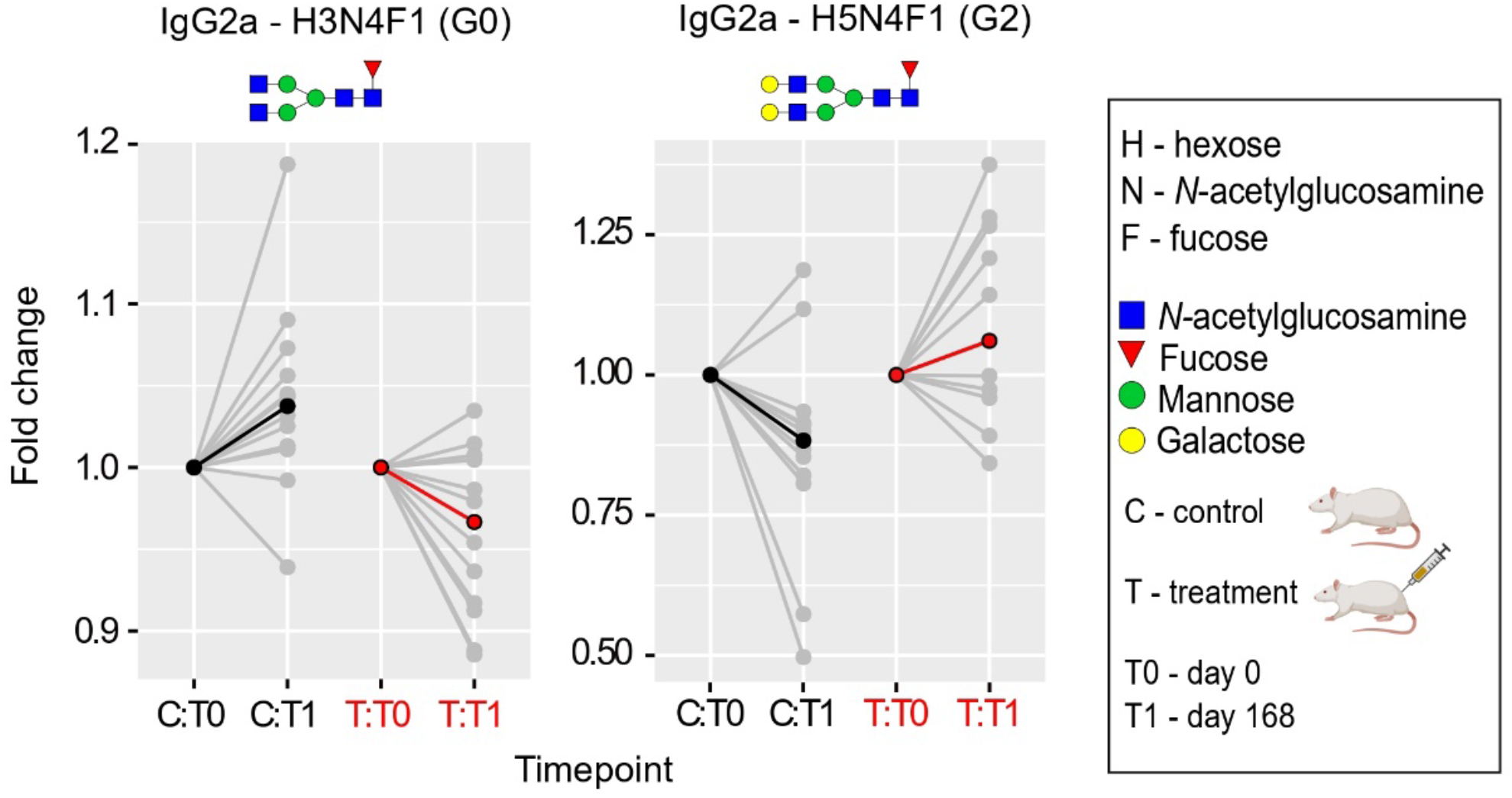
Plasma Fraction Treatment Induces a Shift from Pro-to Anti-Inflammatory Glycan Aging Pattern in Rat IgG2a N-Glycoproteomics. Following the administration of the third E5 dose, rat serum was sampled on the fifteenth day for subclass-specific immunoglobulin G (IgG) Fc N-glycosylation analysis using nano-LC-MS. Notably, a significant (p < 0.05) reduction in the relative abundance of the pro-inflammatory agalactosylated IgG2a glycoform (G0) was recorded. This was accompanied by a simultaneous upsurge in the anti-inflammatory digalactosylated glycoform (G2). The glycoproteomic data was evaluated using a linear mixed-effects model. Median values are emphasized in black (for controls) or red (for treatment) at each time point, with the relative change being normalized to baseline. N-glycan illustrations are provided, utilizing the symbol nomenclature endorsed by the Consortium for Functional Glycomics.

On the fifteenth day after administering the third dose of E5, rat serum was collected for subclass-specific immunoglobulin G (IgG) Fc N-glycosylation analysis using nano-LC-MS. We observed a significant (p < 0.05) decrease in the relative abundance of the pro-inflammatory agalactosylated IgG2a glycoform (G0), which was accompanied by a corresponding surge in the anti-inflammatory digalactosylated glycoform (**Figure 7** and **Supplementary Table S9**). While IgG monogalactosylation (IgG2a-G1) also adheres to this trend in the E5-treated group of rats (Supplementary Table S9), this glycoform exhibits a stronger correlation with specific diseases and general health rather than aging ^32^.

## Discussion

Decades ago, parabiosis experiments revealed the potential of young circulation in benefiting older mice and rats’ organ function. More recent work has begun to leverage this fascinating discovery for medical applications ^5–12, 39, 40^. The early presumption was that this improvement signifies rejuvenation, but it also could represent functional enhancements without altering epigenetic state or age. This ambiguity underlined the question: what is aging?

Our study presents compelling evidence for epigenetic age reversal in rats via the administration of exosome-containing fraction derived from plasma of piglets. Following our initial report in 2020 ^45^, additional validation and cross-species evidence affirmed our results ^43, 46–48^. Mice studies confirmed that young plasma, administered through heterochronic parabiosis, rejuvenated solid organs, reduced epigenetic age by up to 30%, and extended both lifespan and healthspan, with the effect persisting post-separation ^47^.

Given the limitations of mouse models for extensive blood collection, we developed six rat-specific epigenetic clocks using DNA methylation data from thirteen tissue types, aligning with our Mammalian Methylation Consortium’s multi-species clocks ^49^. This research encompassed creating rat-specific clocks, studying plasma fraction treatment effects on epigenetic age, and investigating its impact on aging biomarkers, including a novel rat-specific biomarker for Immunoglobulin G N-glycan aging.

Despite disparate data distribution between species with different lifespans, we managed to establish a human-rat pan-tissue relative age clock, offering biologically significant values by displaying relative biological age within each species. Utilizing a mammalian DNA methylation array profiling 36 thousand conserved probes across various mammalian species was crucial in achieving this.

E5 treatment exhibited great potential, significantly reducing epigenetic ages across multiple rat organs, reduced markers of chronic inflammation and oxidative stress, and increased antioxidant levels. These features are particularly important because oxidative stress and inflammation are two fundamental physiological processes underlying many different types of pathologies ^50, 51^. Interestingly, the effect of E5 on the brain is modest. This is possibly due to slower tissue turnover ^52, 53^, but a more certain answer can only be found when a better understanding of the mechanism of epigenetic aging is known. Our current understanding of this mechanism points to a close relationship with development and homeostatic body maintenance post-maturity ^27, 53–59^. Regardless of the reason, maze test performance demonstrated improved brain function of E5-treated rats. Whether this reflects healthier brain function due to overall physical improvements or actual molecular brain rejuvenation remains to be explored.

The mechanism of reversing glycan aging is not entirely clear yet, but our previous studies suggested a potential involvement of IL-6 and TNF-α ^50, 51^. The observed reduction in these concentrations, coupled with the reversal of pro-inflammatory IgG glycosylation pattern following treatment, supports the rejuvenating impact of E5.

We found the plasma fraction treatment consistently effective in both male and female rats, drastically reducing the epigenetic age of multiple rat tissues. This suggests a potential paradigm shift in healthcare; rather than treating diseases individually, rejuvenation may systemically reduce disease onset risk.

Our study showcased E5 treatment’s rejuvenating capacity in old rats with just two series of injections. The systemic rejuvenation of non-blood organs and the reversal of the epigenetic clock suggests a promising strategy for disease prevention. While E5 treatment is yet to undergo human trials, there is room for optimism as umbilical cord plasma concentrate-based treatment has demonstrated promising effects in a phase 1 clinical trial ^48^. In this regard, our use and demonstration of the rejuvenating efficacy of plasma fraction from piglets is pivotal. Not only does this reinforce the universality of the ageing process in different mammalian species, but it also demonstrates that the source of the rejuvenating material can be readily obtained from species other than humans, which will greatly address the issue of supply, economics, and ethics, which would otherwise be highly challenging.

## Materials and Methods

### Materials

In total, we analyzed n=613 rat tissue samples from 13 different tissues/organs (**Supplementary Table S1, Supplementary Table S2, Supplementary Table S3**). Ages ranged from 0.0384 years (i.e., 2 weeks) to 2.3 years (i.e., 120 weeks). The rat data were comprised of a training set (n=503, **Figure 1**, **Supplementary Table S1**), a test set for the E5 treatment (n=76 from multiple tissues, **Figure 2**, **Supplementary Table S3**) and a second test set for the E5 treatment (n=34 from blood, **Supplementary Table S3**).

We used n=503 tissue to train 4 clocks: a pan-tissue clock based on all available tissues, a brain clock based on regions of the whole brain - hippocampus, hypothalamus, neocortex, substantia nigra, cerebellum, and the pituitary gland, a liver clock based on all liver samples, and a blood clock.

To build human-rat clocks, we added n=1366 human tissue samples to the training data. We first trained/developed epigenetic clocks using the training data (n=503 tissues, **Supplementary Table S1**). Next, we evaluated the data in independent test data (n=76 for evaluating the effect of E5 plasma fraction treatment (**Supplementary Table S3**).

### Methods

The rat tissues came from 4 different labs across three countries:(i) India: Yuvan Research in collaboration with School of Pharmacy SVKM’s NMIMS University (K. Singh), (ii) United States: University of Tennessee Health Science Center (H. Chen) and Medical College of Wisconsin (L.C. Solberg Woods), and (iii) Argentina: University of La Plata (R. Goya).

### Rats from Tennessee and Wisconsin

Blood samples (n=48): Male and female heterogeneous stock rats were bred at the Medical College of Wisconsin (Solberg Woods Lab) or University of Tennessee Health Science Center (Hao Chen Lab). Heterogeneous Stock (HS) populations were originally developed by breeding together eight inbred strains, followed by maintaining the colony in a manner that minimizes inbreeding, allowing fine-resolution genetic mapping of a variety of complex traits (*53*). Rats were euthanized at different ages by an overdose of isoflurane (> 5%). Trunk blood was collected immediately and stored at -80 °C until processing. Blood samples were treated with streptokinase (60-80 IU/200 µl blood, overnight incubation at 37 °C) and DNA was extracted using the QiaAmp Blood Mini Kit (Qiagen Cat No./ID: 51304) following manufacturer’s instructions. All procedures were approved by the Institutional Animal Care and Use Committee of the University of Tennessee Health Science Center or the Medical College of Wisconsin and followed the NIH Guide for the Care and Use of Laboratory Animals. Genomic DNA was isolated from tissue samples mostly using Puregene chemistry (Qiagen). DNA from liver was extracted manually and from blood using an automated Autopure LS system (Qiagen). From tissues and clotted blood samples DNA was extracted manually using QiaAmp DNA Blood Midi Kit and the DNeasy Tissue Kit according to manufacturer’s protocol (Qiagen, Valencia, CA). DNA from BA10 was extracted on an automated nucleic acid extraction platform Anaprep (Biochain) using a magnetic bead-based extraction method and Tissue DNA Extraction Kit (AnaPrep).

### Rats from the University of La Plata (R. Goya lab)

Multiple tissues/cell types (adipose, blood, cerebellum, hippocampus, hypothalamus, liver, neocortex, ovaries, pituitary, skin, substantia nigra): Young (3.7 mo., n=11), Late Adults (LA, 8.0 mo., n=9), Middle-Aged (M-A, 15.7 mo., n=6) and Old (25.5 mo., n=14) female Sprague-Dawley (SD) rats, raised in our Institute, were used. Animals were housed in a temperature-controlled room (22 ± 2°C) on a 12:12 h light/dark cycle. Food and water were available *ad libitum*. All experiments with animals were performed in accordance to the Animal Welfare Guidelines of NIH (INIBIOLP’s Animal Welfare Assurance No A5647-01) and approved by our Institutional IACUC (Protocol # P05-02-2017).

#### Tissue sample collection

Before sacrifice by decapitation, rats were weighed, blood was withdrawn from the tail veins with the animals under isoflurane anesthesia and collected in tubes containing 10µl EDTA 0.342 mol/l for 500µl blood. The brain was removed carefully severing the optic and trigeminal nerves and the pituitary stalk (not to tear the pituitary gland), weighed and placed on a cold plate. All brain regions were dissected by a single experimenter (see below). The skull was handed over to a second experimenter in charge of dissecting and weighing the adenohypophysis. The rest of the body was handed to other 2 or 3 experimenters who dissected and collected whole ovaries, a sample of liver tissue, adipose tissue and skin tissue from the distal portion of tails.

#### Brain region dissection

Prefrontal cortex, hippocampus, hypothalamus, substantia nigra and cerebellum were rapidly dissected on a cold platform to avoid tissue degradation. After dissection, each tissue sample was immediately placed in a 1.5ml tube and momentarily immersed in liquid nitrogen. The brain dissection protocol was as follows. First a frontal coronal cut was made to discard the olfactory bulb, then the cerebellum was detached from the brain and from the medulla oblongata using forceps. To isolate the medial basal hypothalamus (MBH), brains were placed ventral side up and a second coronal cut was made at the center of the median eminence (−3,6 mm referred to bregma). Part of the MBH was taken from the anterior block of the brain and the other part from the posterior block in both cases employing forceps. The hippocampus was dissected from cortex in both hemispheres using forceps. This procedure was also performed on the anterior and posterior blocks, alternatively placing the brain caudal side up and rostral side up. To dissect the substantia nigra, in each hemisphere a 1-mm thick section of tissue was removed from the posterior part of the brain (−4,6 mm referred to bregma) using forceps. Finally, the anterior block was placed dorsal side up, to separate prefrontal cortex. With a sharp scalpel, a cut was made 2 mm from the longitudinal fissure, and another cut was made 5 mm from it. Additionally, two perpendicular cuts were made, 3 mm and 6 mm from the most rostral point, obtaining a 9 mm^2^ block of prefrontal cortex. This procedure was performed in both hemispheres and the two prefrontal regions collected in a code-labeled tube.

#### Anterior pituitary

Using forceps the dura matter that covers gland was removed leaving the organ free on the sella turcica. The neural lobe was carefully separated from the anterior pituitary (AP) which was then carefully lifted with fine curved tip forceps pointing upwards. It was rapidly weighed, then put in a tube and placed momentarily in liquid nitrogen.

#### Ovaries

The genital apparatus was dissected by cutting the mesentery to isolate the uterine horns, the tubular oviduct, the ovaries and the junction between the anus/rectum and the vulva/vagina, leaving the unit of the sexual organs and the urinary bladder isolated. The ovaries were carefully separated from the oviducts; the fat around the ovaries was also removed. Both gonads were placed in a single eppendorf tube and momentarily placed in liquid nitrogen.

#### Liver

Liver tissue extraction was made by cutting a piece of the median lobe (0.5 cm x 0.5 cm). Tissue was placed in a tube and momentarily stored immersed in liquid nitrogen.

#### Adipose tissue

Adipose tissue samples were obtained from the fatty tissue of the small intestine.

#### Tail skin

For skin tissue, 5 cm of a distal tail portion were cut with scissors. Skin was separated and hair removed using scalpel. Tissue was placed in a tube and stored as described for other tissues.

DNA was extracted from blood on an automated nucleic acid extraction platform called QiaSymphony (Qiagen) with a magnetic bead-based extraction kit, QIAsymphony DNA Midi Kit (Qiagen). DNA was extracted from tissue on an automated nucleic acid extraction platform called Anaprep (Biochain) with a magnetic bead-based extraction kit, Tissue DNA Extraction Kit (Biochain). DNA from brain regions was extracted using an automated nucleic acid extraction platform called QIAcube HT (Qiagen) with a column-based extraction kit, QIAamp 96 DNA QIAcube HT Kit (Qiagen).

### Rats from Yuvan Research Group

The Sprague Dawley rats were procured from the National Institute of Bioscience, Pune, India. Animals were housed in the animal house facility of School of Pharmacy, SVKM’s NMIMS University, Mumbai during the study under standard conditions (12:12 h light: dark cycles, 55-70% of relative humidity) at 22±2°C temperature with free access to water and standard pellet feed (Nutrimix Std-1020, Nutrivet Life Sciences, India). The animals were acclimatized to laboratory environment for seven days before initiation of the study. The experimental protocol was approved by the Institutional Animal Ethics Committee. The approval number is CPCSEA/IAEC/P-75/2018.

Sprague Dawley rats of both sexes were used, from which blood, whole brain, heart and liver were harvested. Three batches or samples were prepared.

The first batch was intended for training the epigenetic clock: n=42 blood samples, n=18 whole brain, n=18 heart, n=18 liver samples.

The second batch of test set involved n=76 tissue samples from male rats (n=22 blood, n=18 liver, n=18 heart, n=18 hypothalamus). The test data were used to evaluate the effect of the treatment in 3 conditions: young (30 weeks old) and treated old samples (109 weeks old), and untreated old samples (again 109 weeks old). We evaluated 4 sources of DNA: blood (n=18), liver (n=18), heart (n=18) and hypothalamus (n=18).

Rats were euthanized at different ages by an overdose of isoflurane (> 5%).

The third batch of blood data was collected from both male (n=7) and female (n=10) Sprague Dawley rats. Of these, n=9 animals received E5, while n=8 animals only received a saline injection (control). We collected two blood draws from each animal: at baseline (26 months old) and 2 weeks after treatment.

Trunk blood was collected immediately and stored at -80°C until processing. 100 µl blood sample was treated with 20 µl Proteinase K and then the volume was adjusted to 220 µl with Phosphate Buffer Saline (PBS) in 1.5 ml or 2 ml microcentrifuge tube. 200 µl buffer AL was mixed thoroughly to this mixture by vortexing, and incubated at 56°C for 10 min. Then 200 µl ethanol (96–100%) was added to the sample and mix thoroughly by vortexing and DNA were extracted using the Qiagen DNeasy blood and tissue kit, Qiagen Cat No./ID: 69504 following manufacturer’s instructions. The study protocol was approved through Institutional Animal Ethics Committee (approval no. CPCSEA/IAEC/P-6/2018) which was formed in accordance with the norms of the Committee for the Purpose of Control and Supervision of Experiments on Animals (CPCSEA), Government of India and complied with standard guidelines on handling of experimental animals.

### Rats used in the glycan study

The rat samples for the IgG glycan aging measurement were from the above-mentioned mixed gender study but there is one notable difference: while the methylation study used samples extracted at day zero and just 15 days into the study, the glycan aging study used samples collected at day zero and at 168th day (15 days after 3rd dose). The date of day zero sample was June 17, 2022 while the date of the second time point/sample was December 2, 2022. We also used 6 young controls in the glycan aging study. Thus, we profiled serum IgG N-glycome from 6 female treated, 6 female control, 6 male treated, 6 male control, and 6 young control. The older rats were all born between 17th to 19th April 2020. Young control rats were born between February and April 2022.

### Human tissue samples

To build the human-rat clock, we analyzed previously generated methylation data from n=1366 human tissue samples (adipose, blood, bone marrow, dermis, epidermis, heart, keratinocytes, fibroblasts, kidney, liver, lung, lymph node, muscle, pituitary, skin, spleen) from individuals whose ages ranged from 0 to 93. The data come from our study of primates ^60^. Tissue and organ samples from the National NeuroAIDS Tissue Consortium ^61^. Blood samples from the Cape Town Adolescent Antiretroviral Cohort study ^62^. Skin and other primary cells provided by Kenneth Raj ^54^, Ethics approval (IRB#15-001454, IRB#16-000471, IRB#18-000315, IRB#16-002028).

### Plasma fraction E5

The plasma fraction termed “E5” was developed by Harold Katcher and Akshay Sanghavi at Yuvan Research. This fraction was derived from the platelet-free plasma of young pigs (6-7 months old), an age at which pigs are typically slaughtered by farmers. The selection of this age, immediately post-puberty, is reflective of the mammalian youthful homeostatic peak, with the anticipation that this would be represented in the regulatory cargo circulating within the secretome ^63^. This period marks the beginning of major aging-related changes globally ^64^. The E5 treatment was formulated using plasma from pasture-fed, hormone-free tribal pigs (of the Yorkshire breed) in India. We have found it more complex to create E5 from porcine plasma obtained from pig farms in America.

To this platelet-free plasma, an equal volume of PEG solution in 0.5 Molar NaCl was added, and the combined mixture was incubated for 8 to 12 hours at 4°C. After the stipulated time, the solution was centrifuged at 1000xg for five minutes at 4°C, and the supernatant was discarded. The sediment collected at the bottom of the tubes was transferred into a vessel and mixed with enough physiological saline buffer to make a suspension. The suspension obtained in the above step was subjected to size-exclusion chromatography. The fractions collected from size exclusion chromatography were recombined and concentrated. The concentrate was then mixed with sterile physiological saline solution and injected into rats through the tail vein.

In the following we will provide more details on the treatment. Pig blood, sourced from pasture-raised pigs in India, was collected during slaughter via both the carotid artery and heart puncture. The collected blood was treated with acid citrate dextrose to prevent hemolysis and platelet activation, which we anticipated might dilute non-rejuvenating factors. We aimed to keep the blood temperature around 35 C to avert platelet aggregation. Approximately 1.5 liters of blood was procured per 60 kg animal.

The blood was subjected to centrifugation at room temperature (about 30 C in Mumbai) at 500 x g for 15 minutes. This allowed separation of the blood cells and platelet-rich plasma. Post-centrifugation, the plasma fraction was carefully collected using a pipet, ensuring no loss of larger ectosomes and exosomes. This plasma was further centrifuged at 2,500 x g for 25 minutes to pellet remaining platelets, and the supernatant platelet-poor plasma was stored at 4 degrees C.

Considering the requirement for large volumes of plasma, we employed a modified Extra-PEG method for exosome purification. This method involved an overnight precipitation of a plasma-PEG solution at 4 C, followed by a low-speed centrifugation. Given the observed concentration-dependency of nanoparticle numbers and protein-to-exosome ratios, we opted to use a final PEG concentration of 12%.

To eliminate protein aggregates and fibrin filaments, the redissolved PEG precipitate was passed through a Sephadex G100 column. The resulting fractions were pooled, concentrated via overnight dialysis, and stored appropriately depending on the intended use.

### Experimental procedure

Although transfusion technologies for humans are well-developed and safe, transfusion of small animals is still at the infancy stage of development, requiring state-of-the-art techniques and remains challenging. The necessary dose of E5 was prepared and injected intravenously using sterile saline as vehicle.

Several assumptions underpinned the dosing for old rats (18 to 24 months old). These included estimations of blood mass as a percentage of body mass, the half-volume of plasma, the presence of pro-aging factors in old rat plasma, equivalence in concentration of anti-aging factors between young pig and rat blood, and preservation efficacy of the PEG process. Based on these assumptions, and in consideration of concentration-dependency of exosome rejuvenation, the dosing was calculated such that the delivered pig anti-aging components would quadruple the concentration that would be present in young rat blood. This amounted to 1.43 grams of solid precipitate per 500 grams rat, administered as four injections every other day via tail veins.

The calculated doses were administered intravenously to the animals of old treated group; 4 injections every alternate day for 8 days, and a second dosing starting from the 95^th^ day consisting of 4 injections every alternate day for 8 days, as shown in Supplementary Figure 4. Similar amount of sterile saline solution (placebo) was administered to the animals of old control group. Body weight, food and water intake of the animals were monitored at each time point. Cognitive abilities of animals were evaluated using Barnes Maze apparatus (spanning a week of training) 1, 2, 3 and 4 months from the start of the 1^st^ series of injections. Blood samples were withdrawn at predetermined time intervals by retro orbital plexus during the treatment for hematological evaluation. Serum was separated from the blood samples of each animal and evaluated for biochemical parameters. Plasma was separated from the blood samples of each animal and was used for evaluation of inflammatory markers i.e., TNF-α and IL-6. Animals were sacrificed from each group at 155^th^ day of treatment and vital organs (brain, heart, lung and liver) of these animals were harvested for testing of oxidative stress biomarkers, level of Nrf2, histopathological and immunohistochemistry studies.

### End-point evaluations

#### Body Weight

Body weights of rats were recorded before the initiation of treatment protocol and then 30, 60, 90,120 and 155^th^ day.

#### Grip strength

A grip strength meter was used to measure forelimb grip strength which represents the muscle strength of animals. Briefly, as rat grasped the bar of muscle strength meter, the peak pull force was recorded on a digital force transducer. Tension was recorded at the time the rat released its forepaws from the bar. Six consecutive measurements were taken per day at intervals of one-minute.

#### Barnes Maze Learning Ability

The Barnes maze platform (91 cm diameter, elevated 90 cm from the floor) consisted of 20 holes (each 5 cm in diameter). All holes were blocked except for one target hole that led to a recessed escape box. Spatial cues, bright light, and white noise were used to motivate the rat to find the escape during each session. For the adaptation phase, each rat explored the platform for 60 s. Any rat that did not find the escape box was guided to it and remained there for 90 s. For the acquisition phase, each trial followed the same protocol, with the goal to train each rat to find the target and enter the escape box within 180 s. Rat remained in the box for an additional 60 s. Four trials per day, approximately 15 min apart, was performed for 6 consecutive days. A Barnes maze apparatus used to determine the learning ability of the animals upon treatment. Performed in each month of experiment.

#### Hematological tests

Blood was collected from the retro-orbital plexus using heparinized capillary tubes before the treatment and on 60^th^ and 155^th^ day of the experiment. One portion of the blood was kept in plain bottles from which serum was collected and stored for biochemical analysis. The other portion was directly subjected for the estimation of various hematological parameters using standard instruments. The levels of hemoglobin (Hb), red blood cell count (RBC), packed cell volume (PCV), mean corpuscular volume (MCV), mean corpuscular hemoglobin (MCH), mean corpuscular hemoglobin concentration (MCHC) and platelets were analyzed in the blood samples in all experimental groups.

#### Biochemical test

Blood samples were collected from the retro-orbital plexus using heparinized capillary tubes before the treatment and on 30^th^, 60^th^, 90^th^, 125^th^ and 155^th^ day of the experiment. One portion of the blood was kept in plain bottles from which serum was collected and stored for biochemical analysis. Further, the levels of serum glutamate pyruvate transaminase (S.G.P.T-IU/L) were carried out by kinetic method recommended by International Federation of Clinical Chemistry (IFCC). All the tests were performed with commercially available diagnostic kits (Erba Mannheim, Germany on Erba Mannheim biochemistry semi auto analyzer). Kidney function tests such as determination of serum creatinine (mg/dl) and uric acid (mg/dl) levels were done according to modified Jaffe’s reaction with commercially available diagnostic kits (Erba Mannheim, Germany on Erba Mannheim biochemistry semi auto analyzer). Blood glucose level (Random) (mg/dl) (Gaikwad et al.,2015), Total protein (g/dl), Total Bilirubin (mg/dl), Direct Bilirubin (mg/dl), Triglyceride (mg/dl), HDL (mg/dl), Cholesterol (mg/dl), Albumin (g/dl) (Erba Mannheim) were determined.

#### Oxidative stress evaluation

At the end of experiment, brain, heart, lung and liver were isolated and 10% tissue homogenate was prepared in ice-cold 50 mM PBS (pH 7.4) by using homogenizer followed by sonication for 5 min. The homogenate was centrifuged at 2000 g for 20 min at 4°C and the aliquots of the supernatant were collected and stored at -20° C up to further evaluation.

#### Estimation of extent of lipid peroxidation (LPO) (Malondialdehyde (MDA)

The brain, heart, lung and liver tissue homogenate samples were treated with 1 % phosphoric acid solution and aqueous solution of 0.6 % thiobarbituric acid. The reaction mixture was heated at 80⁰C for 45 minutes, cooled in an ice bath and extracted with 4.0 ml of N-butanol. The n-butanol layer was separated, and the absorbance of the pink complex formed was estimated at 532 nm as an indicator of extend of lipid peroxidation.

#### Estimation of reduced glutathione (GSH)

The GSH content in the brain, heart, lung and liver tissue homogenate was determined by treating the homogenate with sulfhydryl reagent 5,5’-dithio-bis (2-nitrobenzoic acid) (DTNB) method. Briefly, 20 μl of tissue homogenate was treated with 180 μl of 1mM DTNB solution at room temperature. The optical density of resulting yellow color was measured at 412 nm using a microplate spectrophotometer (Powerwave XS, Biotek, USA).

#### Determination of the catalase activity

The brain, heart, lung and liver tissue homogenate (20 μl) were added to 1 ml of 10mM H_2_O_2_ solution in the quartz cuvette. The reduction in optical density of this mixture was measured by using spectrophotometer in UV mode at 240nm. Rate of decrease in the optical density across three minutes from the addition of heart homogenate was taken as an indicator of the catalase activity present in the homogenate.

#### Estimation of superoxide dismutase (SOD) activity

The brain, heart, lung and liver tissue homogenate (20 μl) were added to a mixture of 20 μl of 500 mM of Na_2_CO_3_, 2 ml of 0.3 % Triton X-100, 20 μl of 1.0 mM of EDTA, 5 ml of 10 mM of hydroxylamine and 178 ml of distilled water. To this mixture, 20 μl of 240 μM of NBT was added. The optical density of this mixture was measured at 560 nm in kinetic mode for 3 minutes at one-minute intervals. The rate increase in the optical density was determined as indicator of the SOD activity.

#### Nrf2 concentration in vital organs

Nrf2 was estimated in brain, heart, lung and liver homogenates using Nrf2 ELISA kit (Kinesis Dx, USA). Organ was removed and homogenate was prepared and kept at -20°C until the execution of assay. The Nrf2 level was determined by using kit according to the manufacturer’s protocol, and the values were calculated from the optical density of samples.

#### Pro-inflammatory cytokines (IL-6 and TNF-α)

The cytokines were estimated in plasma, which was separated from blood of animals and kept at -20^0^C until the execution of assay. The pro-inflammatory cytokine levels including TNF-α and IL-6 were determined by using sandwich ELISA kit (Kinesis Dx, USA), according to the manufacturer’s protocol, and the values were calculated from the optical density.

#### Histopathology of Vital Organs

Brain, heart, spleen, kidney, lung, liver and testis tissues fixed in neutral buffered 10% formalin solution were embedded in paraffin, and serial sections (3 μm thick) were cut using microtome (Leica RM 2125, Germany). The representative sections were stained with hematoxylin and eosin and examined under light microscope (Leica, Germany). The Histopathological sections were screened by a pathologist blinded to the treatments.

#### Oil Red O staining

Cryosections (6 µm thick) were fixed in neutral buffered 10% formalin solution for 10 min. The slides were incubated with freshly prepared Oil Red O working solution for 15 min. Lipid accumulation was digitalized using a microscope.

### DNA methylation profiling

For the first experimental series, we generated DNA methylation data using the mammalian methylation array “HorvathMammalMethylChip40” ^38^. By design, the mammalian methylation array facilitates epigenetic studies across mammalian species (including rats and humans) due to its very high coverage (over thousand X) of highly conserved CpGs in mammals. These 36k CpGs exhibit flanking sequences that are highly conserved across mammals. The subset of species for each probe is provided in the chip manifest file which has been posted on Gene Expression Omnibus. The SeSaMe normalization method was used to define beta values for each probe ^65^.

For the second experimental series, we used the mammalian methylation 320 array platform to generate methylation data from blood. This array combines the CpG content of the mammal 40 array with the Illumina mouse array. The array contains 106K CpGs that map to the genome of the brown rat.

Unsupervised hierarchical clustering was performed to identify outliers and failed arrays, and those were excluded.

### Penalized Regression models

We developed the six different epigenetic clocks for rats by regressing chronological age on the CpGs on the mammalian array. We used all tissues for the pan-tissue clock. We restricted the analysis to blood, liver, and brain tissue for the blood, liver, and brain tissue clocks, respectively. Penalized regression models were created with the R function “glmnet” ^66^. We investigated models produced by both “elastic net” regression (alpha=0.5). The optimal penalty parameters in all cases were determined automatically by using a 10 fold internal cross-validation (cv.glmnet) on the training set. The alpha value for the elastic net regression was set to 0.5 (midpoint between Ridge and Lasso type regression) and was not optimized for model performance. We performed a cross-validation scheme for arriving at unbiased (or at least less biased) estimates of the accuracy of the different DNAm based age estimators. One type consisted of leaving out a single sample (LOOCV) from the regression, predicting an age for that sample, and iterating over all samples.

### Relative age estimation

To introduce biological meaning into age estimates of rats and humans that have very different lifespan; as well as to overcome the inevitable skewing due to unequal distribution of data points from rats and humans across age range, relative age estimation was made using the formula: Relative age= Age/maxLifespan where the maximum lifespan for rats and humans were set to 3.8 years and 122.5 years, respectively.

### Epigenome wide association studies of age and treatment

In our EWAS, we used the same data as those underlying Figure 2.

The normalized beta values were used to fit a least squares linear regression model using the R package limma (v3.50.0) ^67^. The model was adjusted for known variables, including Condition, Sentrix_ID, and methylation array column. To account for unknown confounders, factor analysis was performed using the R package RUVSeq (v1.28.0) ^68^ and two RUVg vectors were added to the model. The defined model was used to identify differentially methylated cytosines in two distinct contrasts: (1) the Age contrast, which aimed to detect differentially methylated loci associated with the aging process, and (2) the Elixir contrast, which compared treated rats with age-matched control rats to determine cytosines that exhibited differential methylation due to the treatment effect. Empirical Bayes statistics were estimated using the eBayes function, and multiple testing was corrected using the Benjamini-Hochberg (BH) method. Probes with an adjusted p-value < 0.05 were considered statistically significant.

The regression models for this study included the following covariates: SID, TreatmentCondition, Sentrix_ID, Col. The full model formula was: ∼ 0 + TreatmentCondition + Sentrix_ID + Col. The null model formula was: ∼ 0 + Sentrix_ID + Col.

Empirical Bayes correction was applied to model fits.

Limma’s contrasts.fit function was used to perform group comparisons. Contrasts used: Elixir = TreatmentConditionOldTreated - TreatmentConditionOldControl

Age = TreatmentConditionOldControl-TreatmentConditionYoungControl

For each dataset and contrast, p values were adjusted with Benjamini-Hochberg correction for multiple testing and those with FDR q < 0.05 were deemed significant.

### Immunoaffinity enrichment of IgG from rat serum

Frozen serum samples were thawed at +4 °C, randomized, and 30 µL was transferred into a 96-well 1-mL collection plate. A total of seven standards prepared by pooling the samples as well as blanks were included to assess the technical variation and possible carryover and cross-contamination, respectively. The IgG was enriched on Pierce^TM^ protein L agarose (Thermo Fisher Scientific Inc.) by downscaling our previously published protocol ^69^. Briefly, the samples were diluted with 270 µL 1x PBS (pH 7.4), 20 µL of protein G beads were added, the plate was sealed with an Easy Pierce heat sealing foil (Thermo Fisher Scientific Inc.) and shaken for 1h on an orbital shaker (800 rpm). Next, the beads were transferred into a 1-mL PP filtration microplate (Agilent, 7 µm frit, cat. no. 202501-100) and washed 3x with 200 µL 1x PBS and 2x 200 µL ultrapure water. The IgG was eluted with 150 µL 0.1 M formic acid (Merck KGaA, Darmstadt) directly into a PCR plate containing 15 µL 1 M ammonium bicarbonate (Acros Organics, Geel, Belgium) solution by centrifugation (4 min, 60 x *g*). The eluate was dried down in a vacuum centrifuge at 40 °C.

### Tryptic glycopeptide preparation and HILIC-SPE

Dry IgG was redissolved in 25 µL 25 mM ammonium bicarbonate and digested with 0.2 µg sequencing grade modified trypsin (Promega Corp., Madison WI, USA) at 37 °C overnight. Tryptic glycopeptides were enriched on Chromabond® zwitterionic HILIC adsorbent (Macherey-Nagel GmbH & Co., Düren, Germany) in a high-throughput manner, dried down in a vacuum centrifuge and stored at -20 °C.

### Nano-liquid chromatography-mass spectrometry and data pre-processing

The rat IgG-Fc N-glycosylation profiling was performed for IgG2a (P20760), IgG2b (P20761) and IgG2c (P20762) subclasses. Dry glycopeptides were redissolved in 30 µL ultrapure water and 5 µL was injected for analysis. The nano-LC-ESI-Q-TOF setup, chromatographic conditions, and MS parameters were thoroughly described in Habazin 2021 PMID: 34118474. MS1 files were first converted to mzXML format using ProteoWizard MSConvert (v. 3.0) and processed with LaCy Tools (v. 1.0.1), Chambers 2012 PMID: 23051804. Relative abundances of IgG N-glycoforms were calculated by normalizing the integrated areas of the extracted ion chromatograms to the total area for each of the three IgG subclasses.

### Glycoproteomic data statistical analysis

Association analyses between chronological age and glycopeptide traits were performed using a general linear model with sex included as additional covariate. Longitudinal analysis of samples through their observation period was performed by implementing a linear mixed-effects model where time was modeled as a fixed effect, the interaction between time and plasma fraction treatment was modeled as a fixed effect, while individual sample ID was modeled as a random intercept. Prior to analyses, glycopeptide variables were all transformed to standard normal distribution (mean = 0, sd = 1) by inverse transformation of ranks to Normality (R package “GenABEL”, function rntransform). Using rank transformed variables in analyses makes estimated effects of different glycopeptides comparable as transformed glycopeptide variables have the same standardized variance. False discovery rate was controlled using Benjamini-Hochberg procedure. Data was analysed and visualized using R programming language (version 4.0.2).

### URLs

## Supporting information

Coefficient values epigenetic clocks for rats

## Acknowledgements

The development of the rat tissue clocks was supported by the Paul G. Allen Frontiers Group (SH) and a grant from Open Philanthropy (SH). The heterogeneous stock rats provided by (HC and LS) were supported by NIH grant DA-037844 (NIDA, HC and LW). RG was supported by grant # MRCF 7-25-19 from the Medical Research Charitable Foundation and the Society for Experimental Gerontological Research, New Zealand (RG). Human tissue sample collection was supported by NIH funding through the NIMH and NINDS Institutes by the following grants: Manhattan HIV Brain Bank (MHBB): U24MH100931; Texas NeuroAIDS Research Center (TNRC): U24MH100930; National Neurological AIDS Bank (NNAB): U24MH100929; California NeuroAIDS Tissue Network (CNTN): U24MH100928 Data Coordinating Center (DCC): U24MH100925. Human blood samples were supported by R21MH107327. The contents are solely the responsibility of the authors and do not necessarily represent the official view of the NNTC or NIH.

## Authors contributions

The plasma fraction treatment was developed by Harold Katcher (HK) in consultation with Akshay Sanghavi (AS). KR, Steve Horvath (StH) and HK drafted the manuscript. All authors helped to edit the article. JZ and StH developed the epigenetic clocks. StH generated the DNA methylation data. JZ, KR, SK, CL, StH carried out statistical analyses and created figures and tables. KS managed the plasma treatment project and carried out the experiments with SK and AgS. RG, HC, CH, LSW, MC-M, ML, PC, TW, AM contributed rat tissue samples. AL and StH contributed human data. SiH, MN, FV and GL carried out immunoglobulin G N-glycosylation analysis and analyzed the glycoproteomic data.

## Supplementary Material

The Supplement contains Supplementary Figures, Supplementary Tables, Supplementary Methods and software code for the rat clocks

**Supplementary Figure 1.**
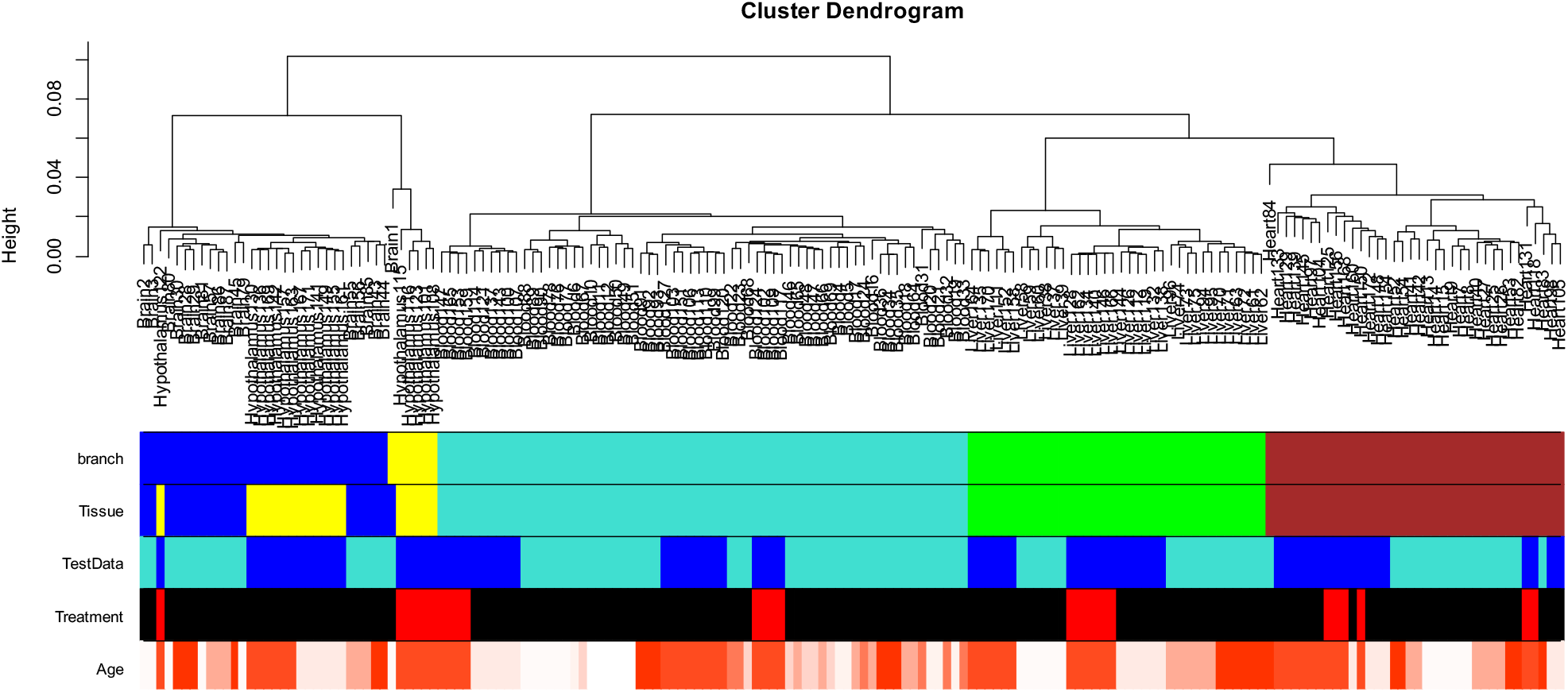
Unsupervised Hierarchical Clustering of Rat Tissue Samples Based on Inter-array Correlations. This figure visualizes the unsupervised hierarchical clustering of rat tissue samples, generated using average linkage clustering based on 1 minus the Pearson correlation as dissimilarity measure (utilizing the R hClust function). Five colorcoded bands below the dendrogram provide additional information: The first band denotes the cluster branch, indicating the grouping of samples. The second band specifies tissue types, represented as blue for brain, yellow for hypothalamus, turquoise for blood, green for liver, and brown for heart. The third band identifies the test data utilized to evaluate the E5 treatment, marked in blue. The fourth band signifies samples that underwent the E5 treatment, represented in red. The fifth band differentiates age groups: white symbolizes young samples, while red stands for older ones. Notably, the branching within the dendrogram closely aligns with tissue types.

**Supplementary Figure 2.**
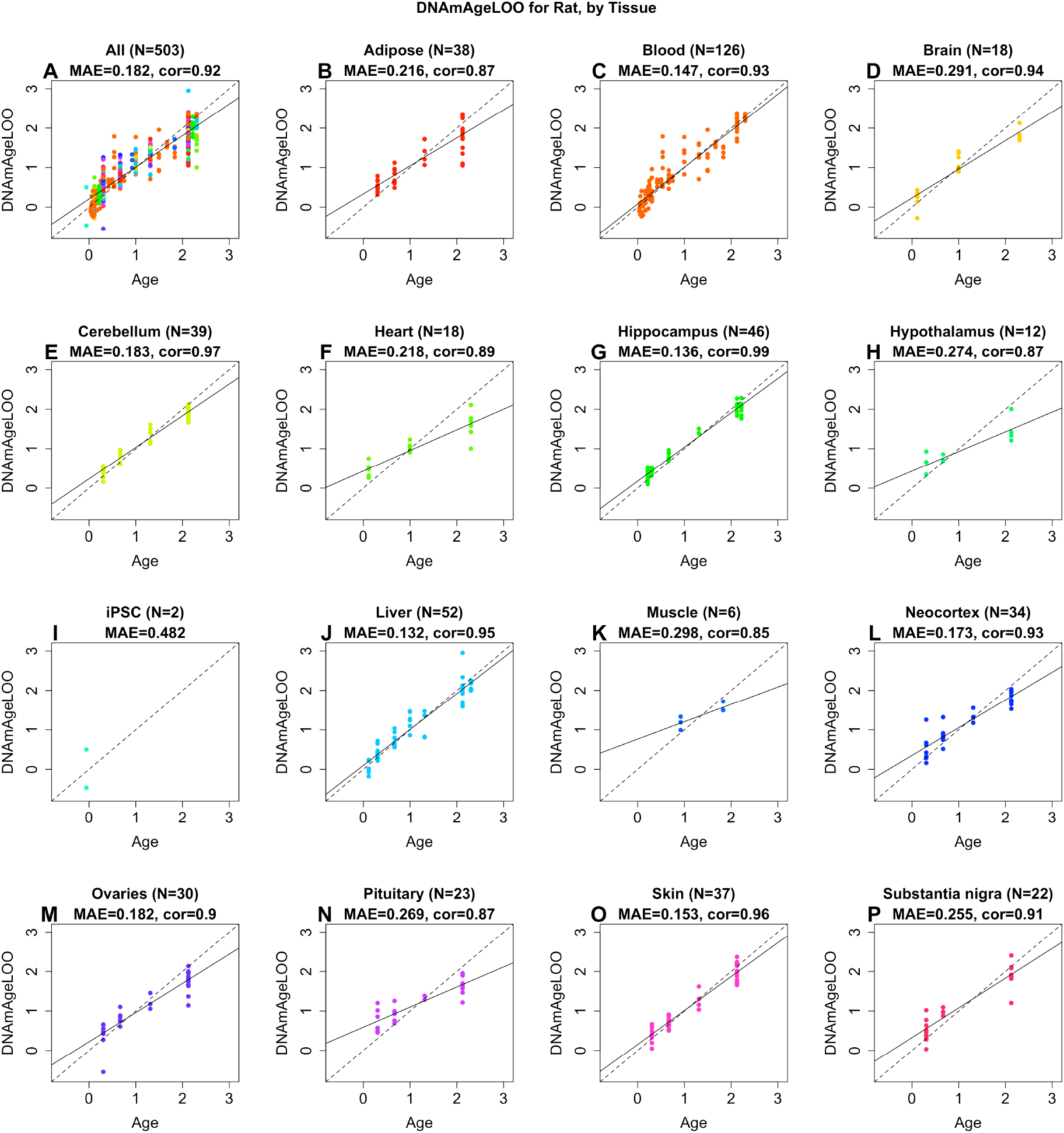
Pan tissue clock for rats applied to different tissues. A) All tissues. B) adipose, C) blood, D) whole brain, E) cerebellum, F) heart, G) hippocampus, H) hypothalamus, I) liver, J) brain neocortex, K) ovaries, L) pituitary, M) skin, N) substantia nigra. The title of each panel reports the tissue, sample size, Pearson correlation coefficient and median absolute deviation (median error). Chronological age (x-axis) versus leave-one-sample-out estimate of age.

**Supplementary Figure 3:**
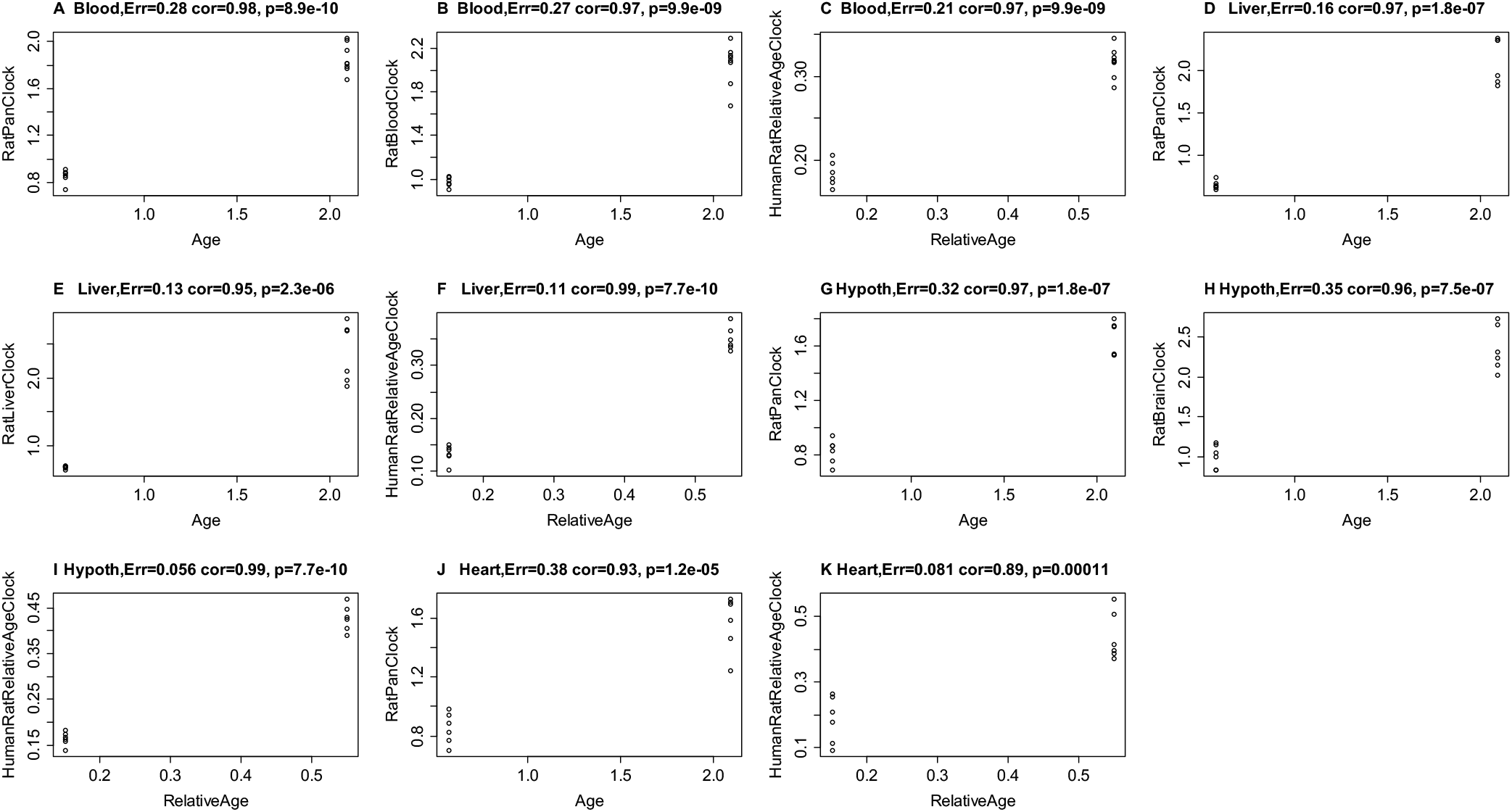
Epigenetic clocks applied to independent test data. The six epigenetic clocks were applied to DNA profiles from un-treated samples from the E5 plasma fraction study. The y-axis reports epigenetic age (in years) as measured by the indicated clocks. The heading reports the tissue type, the correlation between epigenetic and chronological age and the median error (in years). While the age correlations are high, the median errors are sub-optimal, up to 0.38 years (panel J). As such, the final versions of the clocks incorporated un-treated samples from the test data as well (Methods).

**Supplementary Figure 4:**
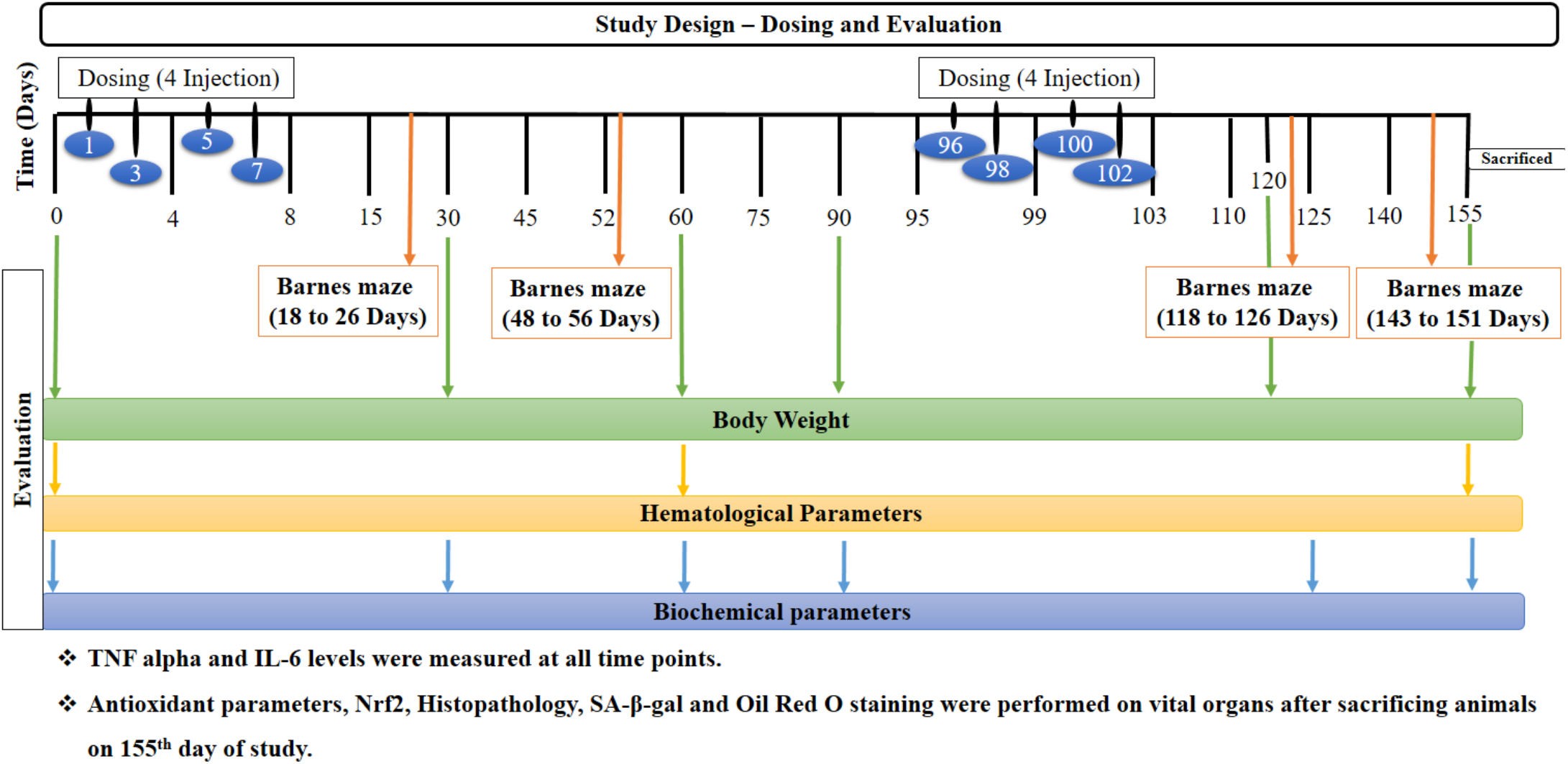
Schematic representation of the study design with timeline indicating plasma fraction treatments as well as cognitive, physical, haematological and biochemical tests.

**Supplementary Figure 5:**
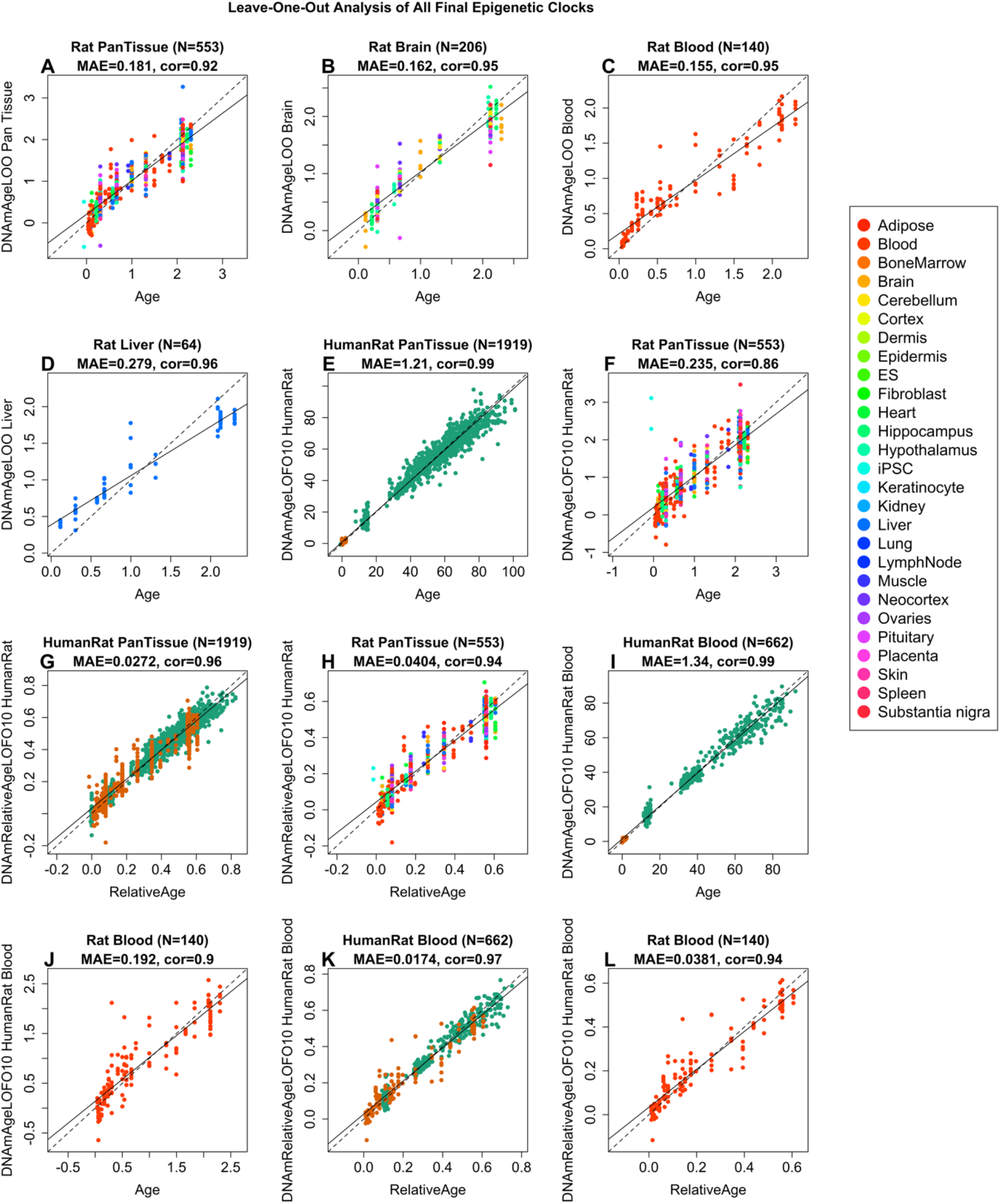
This figure is analogous to Figure 1, but it reports estimates of the predictive accuracy of the final version of the rat clocks. Cross-validation was carried out on an increased number of rat tissues (n=553) by combining the original training data (n=503) with rat tissues from untreated animals of the second test data set.

**Supplementary Figure 6:**
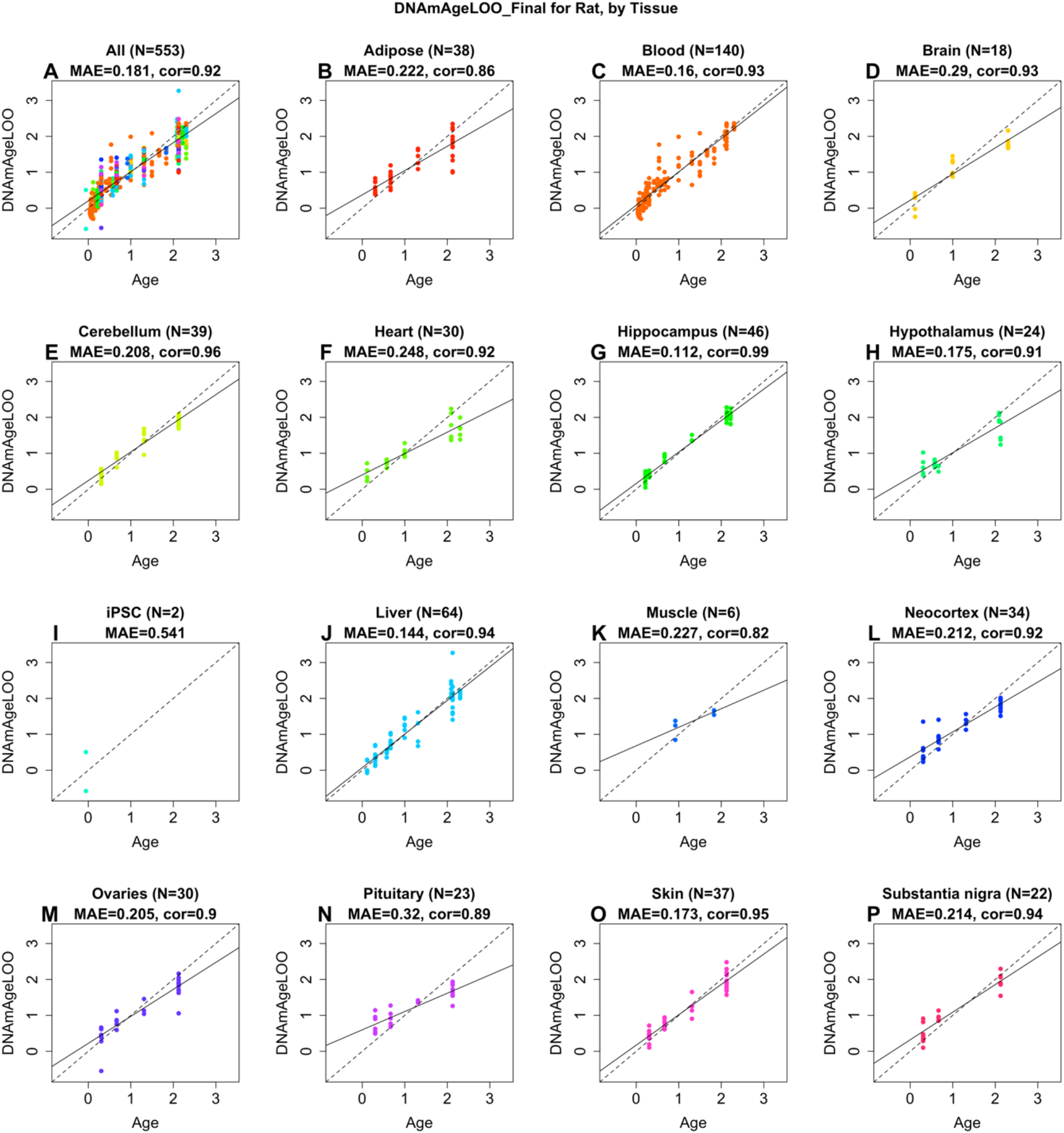
Estimating the accuracy of the final version of the pan tissue rat clock. Each panel corresponds to a different source of DNA in rat tissue. Leave one out cross validation estimates of DNAmAge (y-axis) versus chronological age.

**Supplementary Figure 7.**
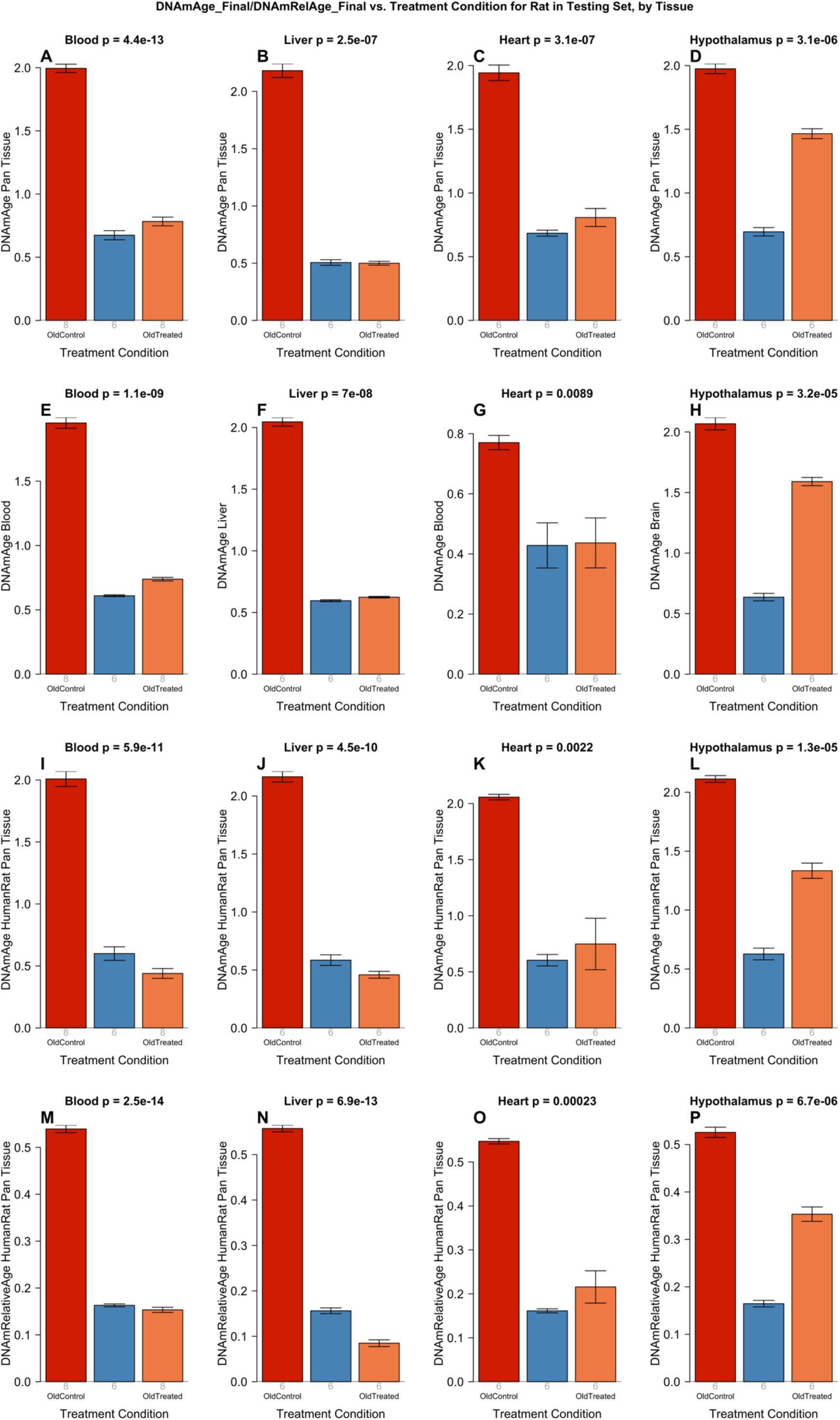
Epigenetic clock analysis of plasma fraction treatment based on the final version of epigenetic clocks. This figure is analogous to Figure 2, but it reports results for the final version of the epigenetic clock based on an increased number of rat tissues (n=567) (resulting from combining the original n=517 original training data with rat tissues from untreated animals of the test data set). Six epigenetic clocks applied to independent test data from four rat tissue type (columns): blood, liver, heart, and hypothalamus. A-D) Rat pan-tissue clock. E) Rat blood clock applied to blood. F) Rat liver clock applied to liver. G) Rat blood clock applied to heart. H) Rat brain clock applied to hypothalamus. I-L) Human-rat clock measure of absolute age. M-P) Human-rat clock measure of relative age defined as age/maximum species lifespan. Each bar-plot reports the mean value and one standard error. Student T-test p values result from a 2-group comparison of old controls (left bar) versus old treated samples (right bar), i.e. the young controls were omitted.

**Supplementary Figure 8 (version 1):**
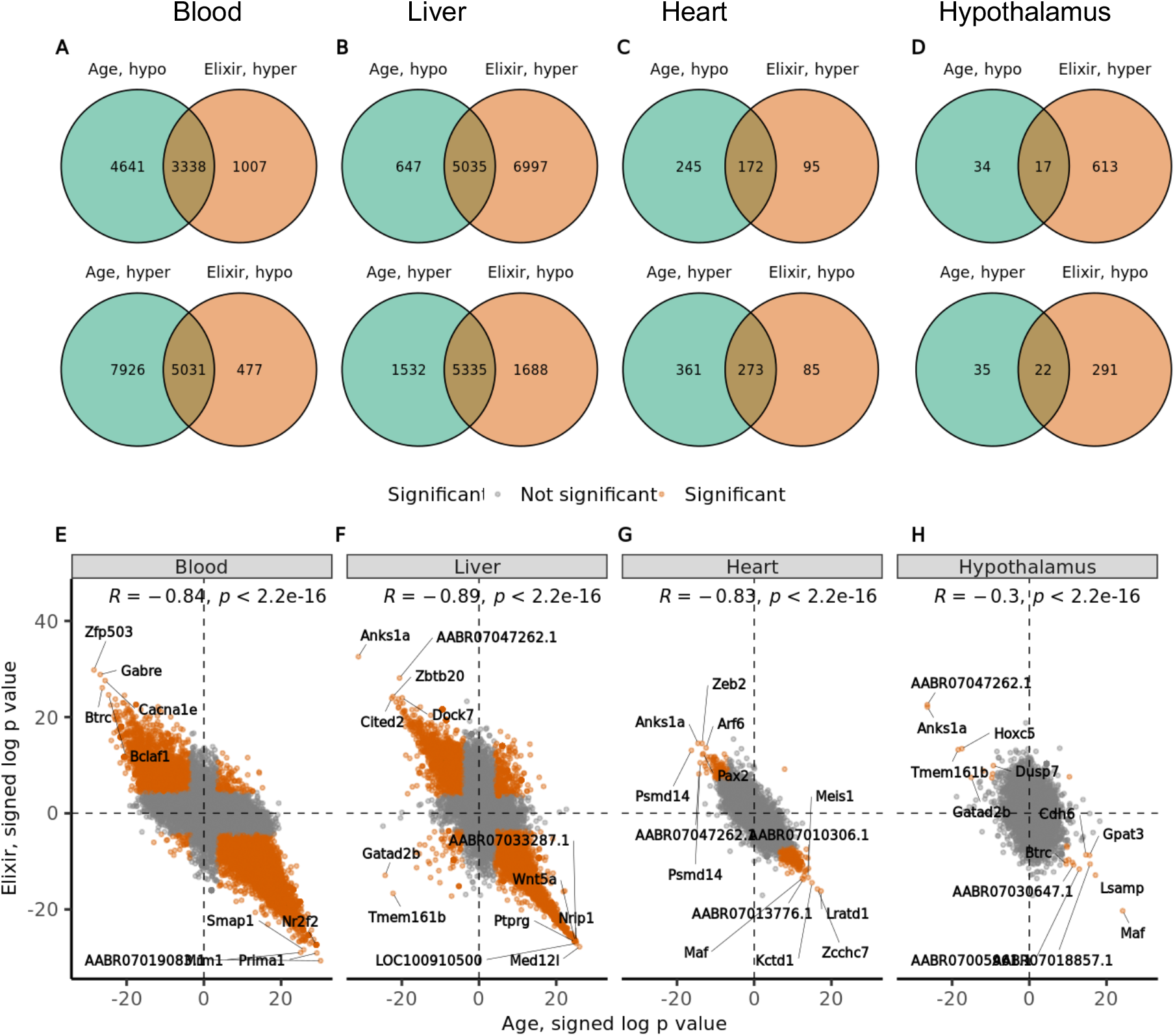
EWAS results for age versus EWAS results of plasma fraction treatment by tissue type. This figure portrays two distinct versions of the EWAS study, differentiated by their respective definitions of aging effects. In Version 1, age effects are determined by comparing old control rats to young ones within the test dataset. This analysis may carry potential bias, as samples from old control animals are employed in assessing both age effects (x-axis) and treatment effects (y-axis). To mitigate this concern, Version 2 of the figure (below) illustrates an alternative approach: age effects are calculated exclusively using independent controls, thus ensuring a more unbiased analysis. The Venn diagrams in version 1 indicate the overlaps of significantly differentially hyper- or hypo-methylated aging and E5 affected CpGs across multiple tissue types: blood (A), liver (B), heart (C), and hypothalamus (D). We used a false discovery rate threshold of 0.05 for both aging and treatment effects (Methods). E-H) Each dot corresponds to a CpG on the mammalian array. Aging effects (x-axis) versus E5 rejuvenation effects (y-axis) in specific rat tissue: blood (E), liver (F), heart (G), and hypothalamus (H). Signed P values were defined as follows: sign(FC) * log(p value). Orange color indicates genes with significant differential methylation in both comparisons. Positive/negative values correspond to gain/loss of methylation associated with the treatment.

**Supplementary Figure 8 (version 2):**
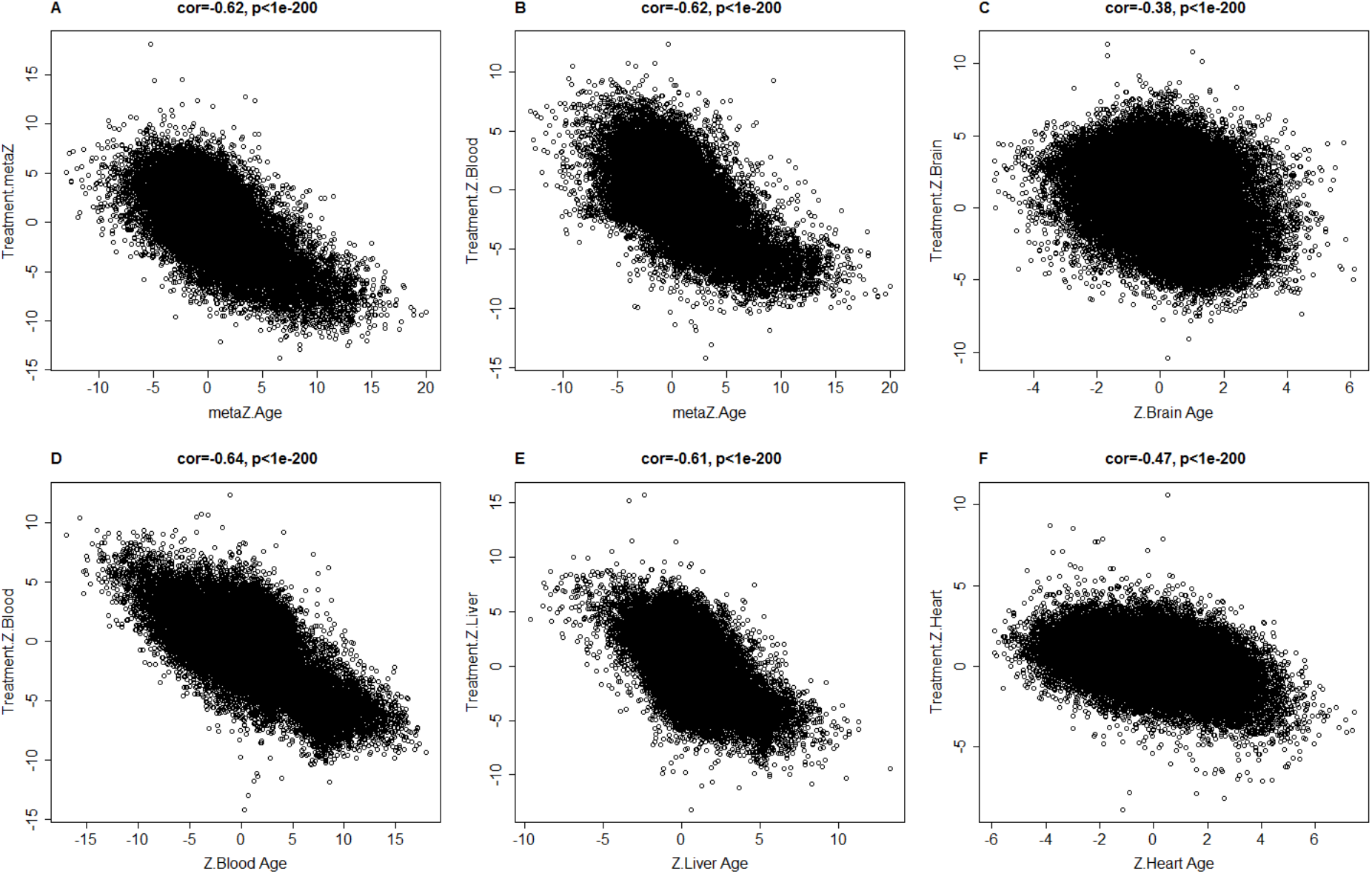
EWAS results for age versus EWAS results of plasma fraction treatment. This version of our EWAS study is less biased than version 1. Each dot corresponds to a CpG on the mammalian array. The x axis reports Z statistics from a correlation test of CpG methylation versus chronological age (R function “standardScreeningNumericTrait” in the WGCNA package). Positive/negative values correspond to positive/negative correlation coefficients with age, respectively. The y-axis reports a Z statistic for the treatment effect. Positive/negative values correspond to gain/loss of methylation associated with the treatment. A-B) The x axis corresponds to a meta-analysis of age effects across all rat tissues (Adipose, Blood, Brain, Cerebellum, Heart, Hippocampus, Hypothalamus, Liver, Neocortex, Ovaries, Pituitary, Skin, Substantia nigra). Age effects in C) Brain, D) Blood, E) Liver, F) Heart. A) Treatment effects across four tissues. Stouffer’s meta-analysis Z statistic across hypothalamus, blood, liver, heart. Treatment effects in individual tissues B) Blood, C) Hypothalamus, D) Blood, E) Liver, F) Heart.

**Supplementary Figure 9A.**
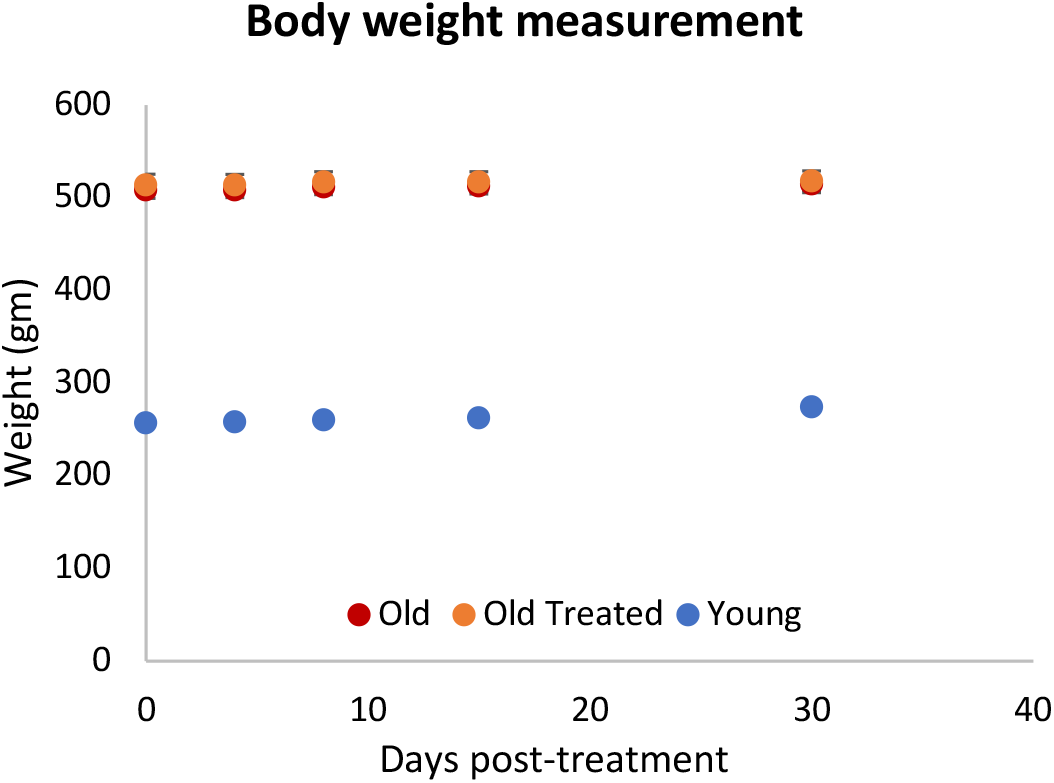
Physical measurements of rats. Weight of rats measured at regular intervals at indicated times during the 155-day experiment. Each group measurement was from 6 rats. The plotted data points represent average values from 6 rats each, with standard error of the mean.

**Supplementary Figure 9B.**
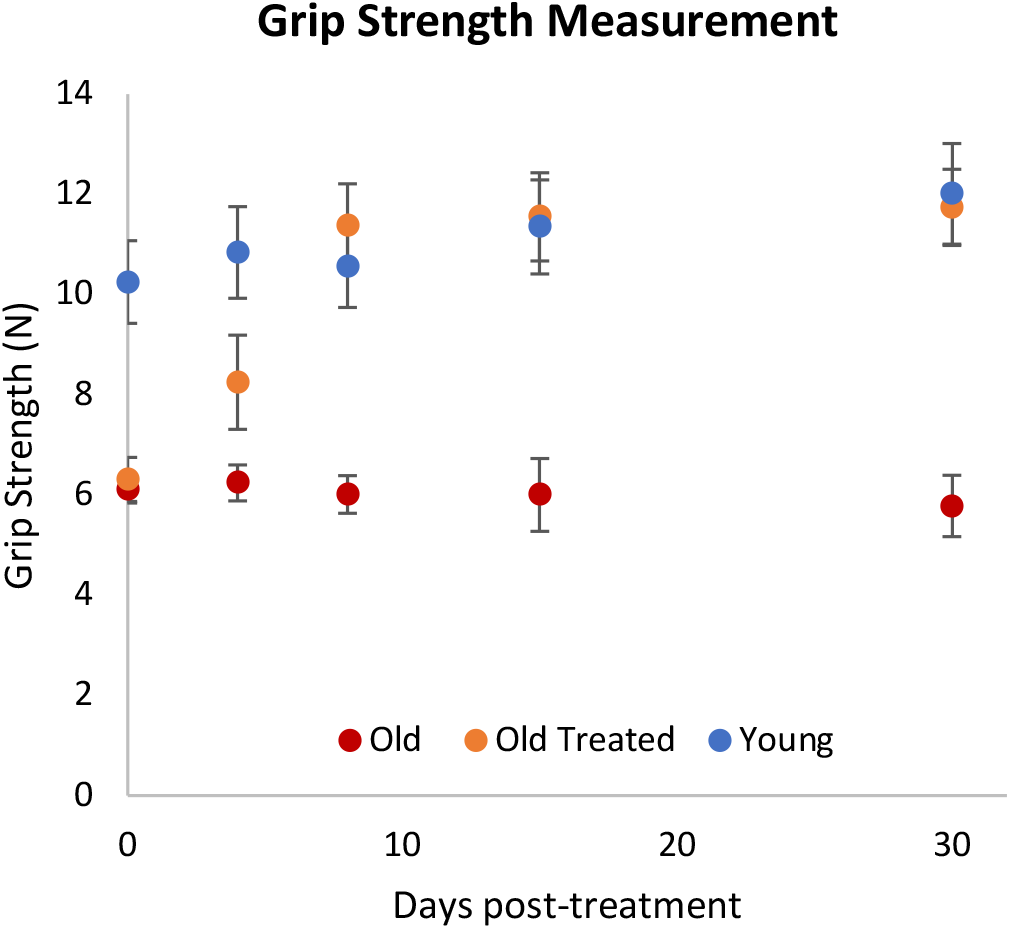
Measurement of grip strength of the indicated rat groups at various times post-treatment. Each group consists of 6 rats. The plotted data points represent average values from 6 rats each, with corresponding 2 standard errors around the mean. Detailed measurements of each parameter are provided in Supplementary Table S5.

**Supplementary Figure 10:**
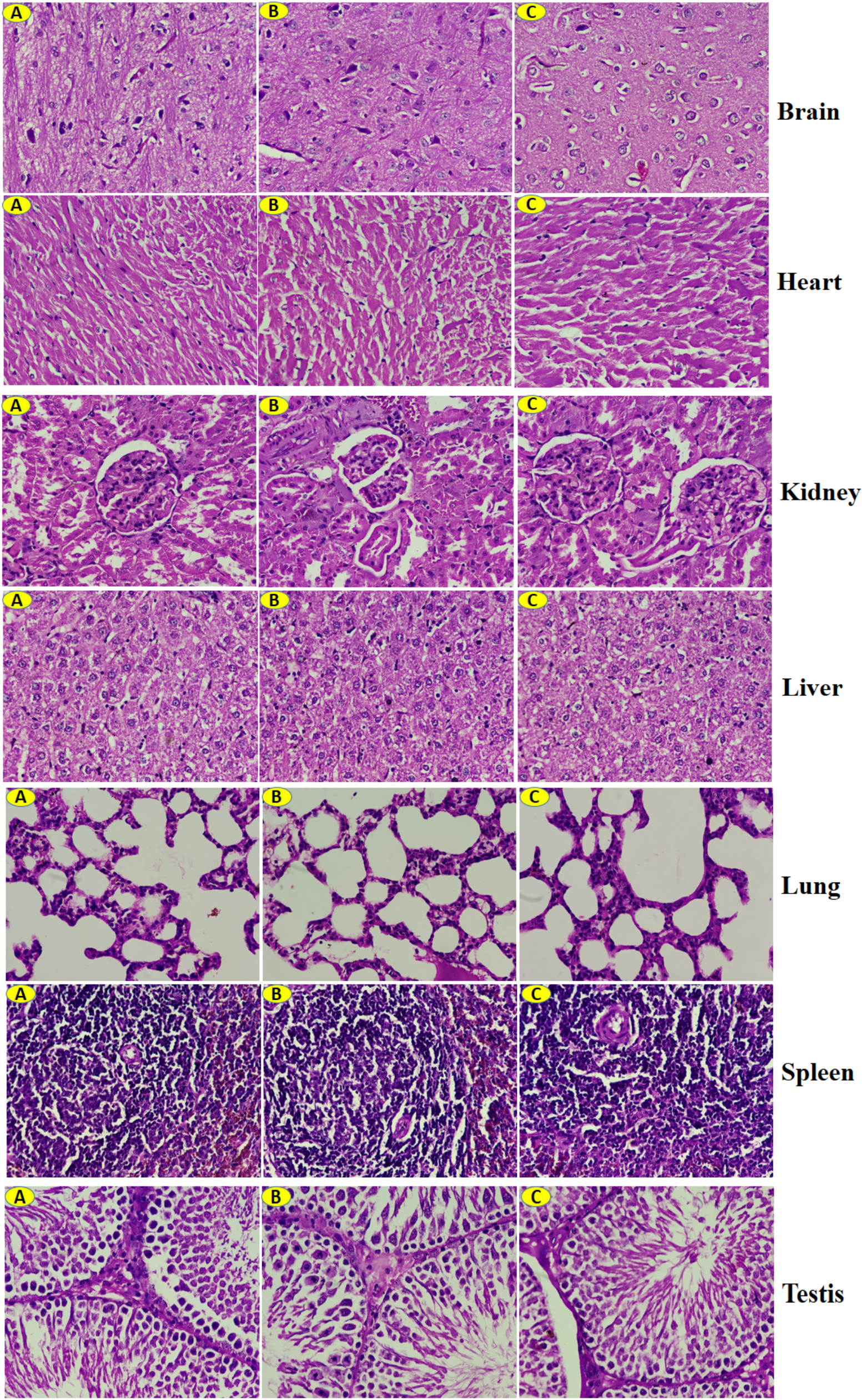
Histological analyses of vital organs and tissues of rats employed in experiment. Images of the left column (A) are from old rats; the right column (C) are images of tissues from young rats while the middle column (B) are images of tissues from old rats treated with plasma fraction. Results of histopathological examination are tabulated in Supplementary Table S6

**Supplementary Figure 11:**
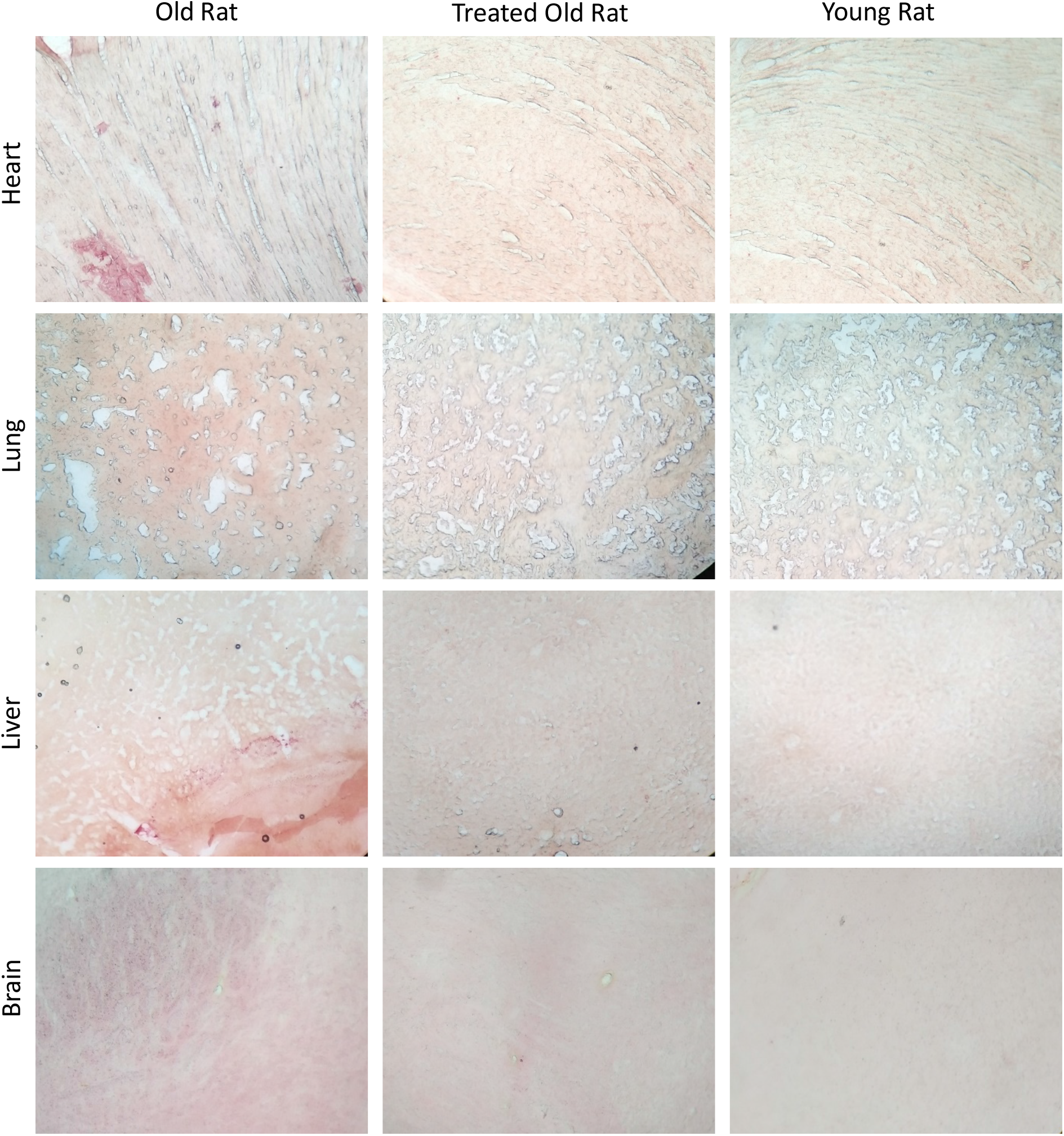
Oil Red O staining. Tissues from old untreated rats, Plasma fraction-treated old rats and young rats were subjected to Oil Red O staining to reveal accumulation of fat in tissues.

**Supplementary Figure 12:**
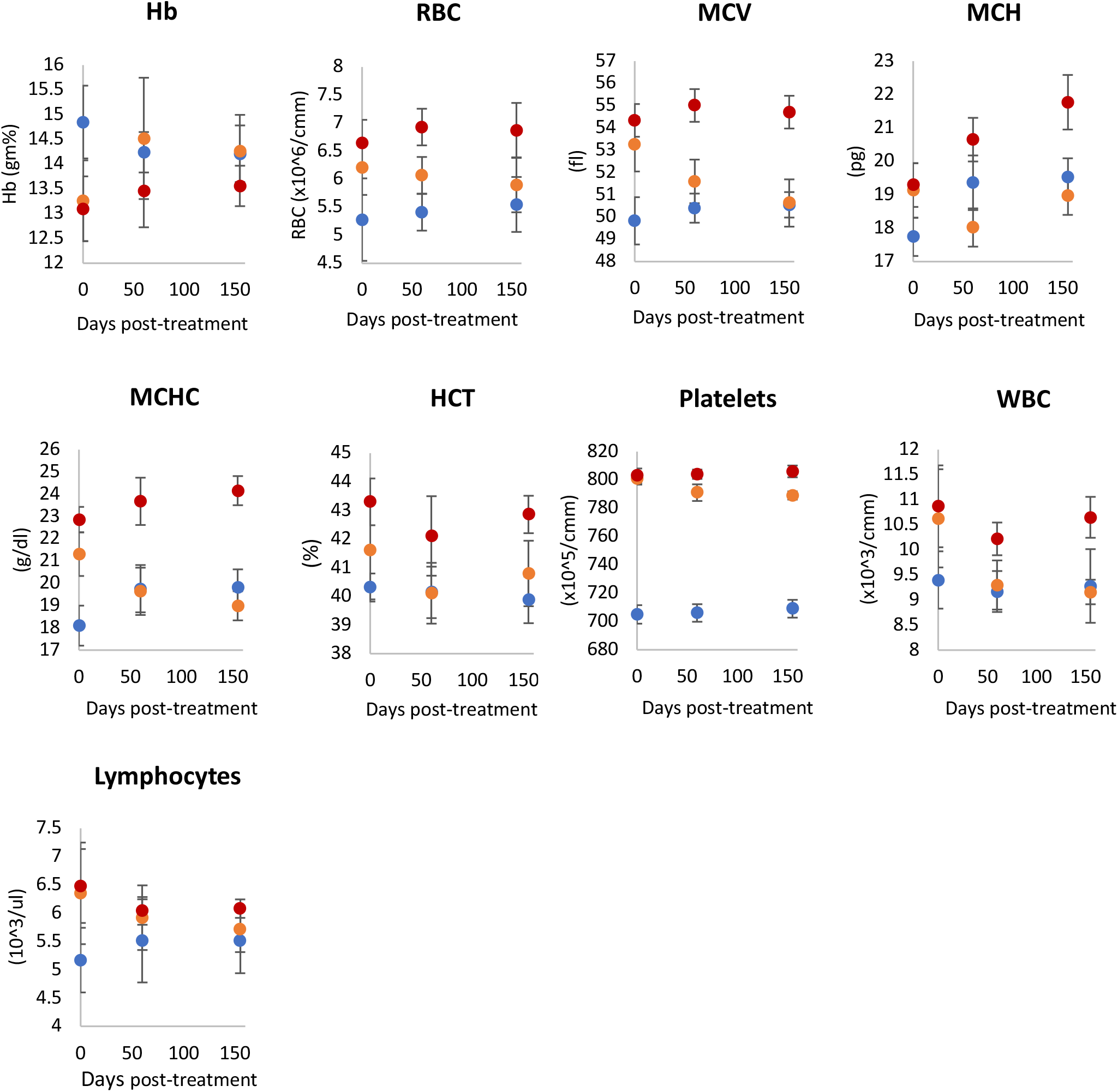
Haematological properties. Effects of plasma fraction treatment on haematological properties of rats at 0, 60 and 155 days from the start of experiment. Hb = haemoglobin, RBC = red blood cell count, MCV = mean corpuscular volume, MCH = mean corpuscular haemoglobin, MCHC = mean corpuscular haemoglobin concentration, HCT = haematocrit and WBC = white blood cells. Red dots represent data points of old rats, orange dots represent treated old rats and blue represents young rats. The plotted data points represent average values from 6 rats each, with 2 standard errors around the mean. Detailed measurements of each parameter are provided in Supplementary Table S7.

**Supplementary Figure 13.**
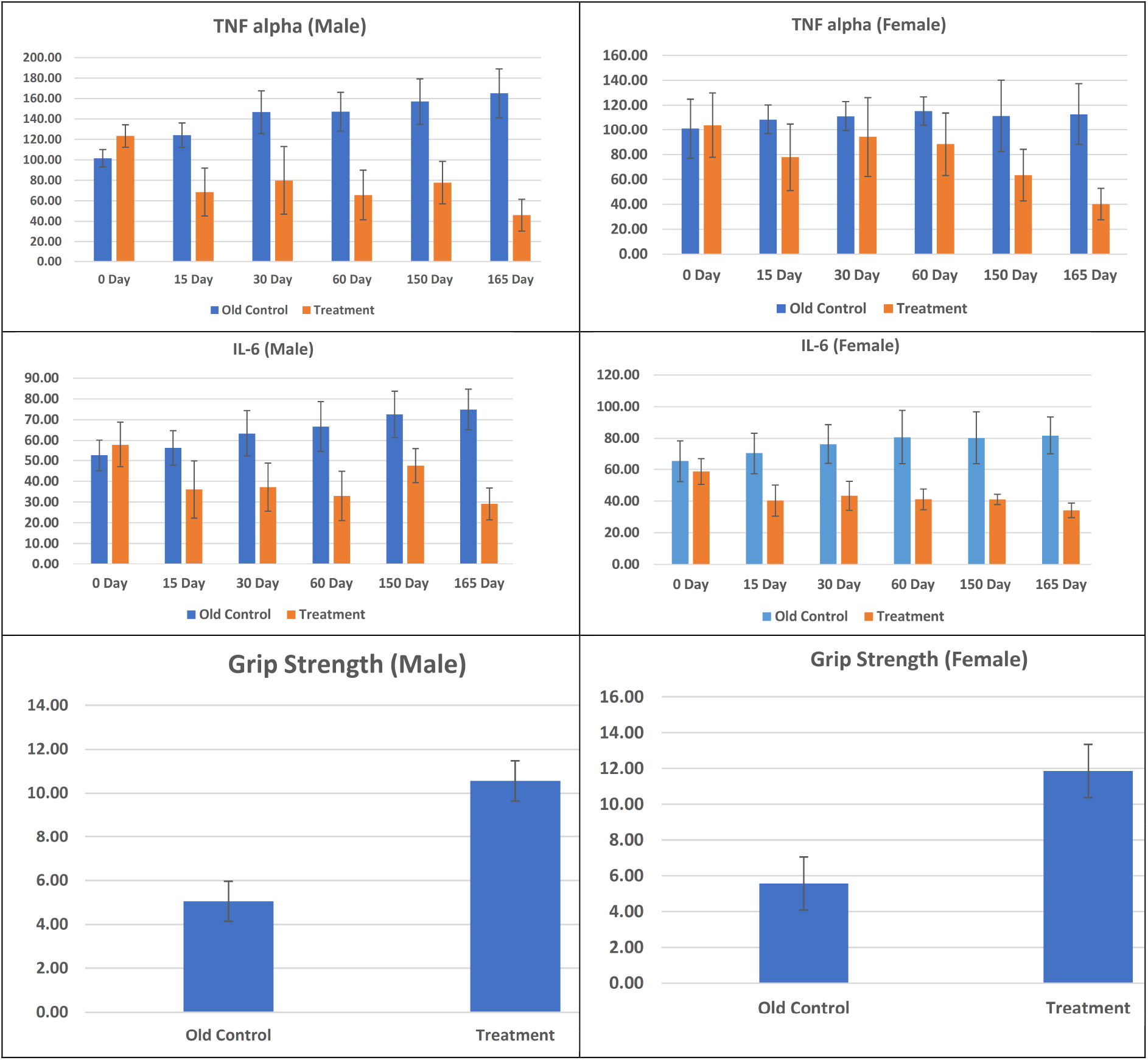
Evaluating the effect of E5 in both male and female Sprague Dawley rats. A,B) TNA alpha (y-axis) and C,D) IL-6 are measured in units of picogram/ml. At baseline, the rats were 26 months old. The study analzed 12 male and 12 female rats. Half of the animals were treated with E5. The controls were treated with saline.

**Supplementary Figure 14.**
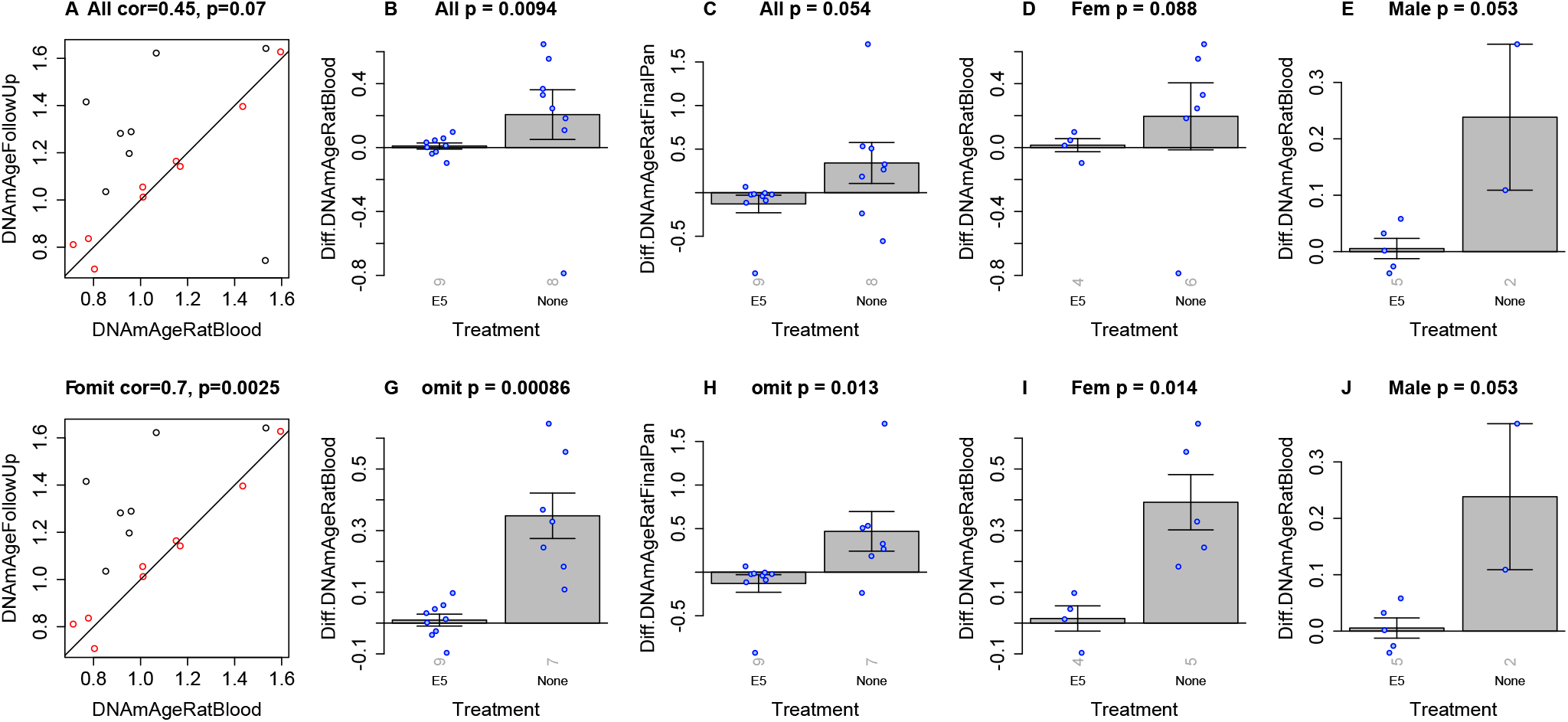
Epigenetic clock analysis in blood of male and female rats. This is an independent validation study of epigenetic clocks that is based on a different set of Sprague Dawley rats of both sexes. A) Final version of the rat clock for blood. Baseline measurement (x-axis) versus follow up measurement (15 days after treatment, y-axis). Points (rats) are colored by treatment: red=treated by E5, black=treated with saline only. B,D,E) Difference between follow up measurement and baseline measurement (y-axis) versus treatment status in B) all rats, D) female rats only, E) male rats only. C) is analogous to B) but uses the pan tissue clock for rats. Panels in the second row (F,G,H,I,J) are analogous to those in the first row but the analysis omitted one control rat (corresponding to the black dot in the lower right of panel A). The title of the bar plots reports the results of a non-parametric group comparison test (unadjusted two-sided p value from the Kruskal Wallis test). The rotated grey numbers underneath each bar reports the group sizes. Each bar plot reports the mean value and one standard error.

## Supplementary Tables

**Supplementary Table S1.**
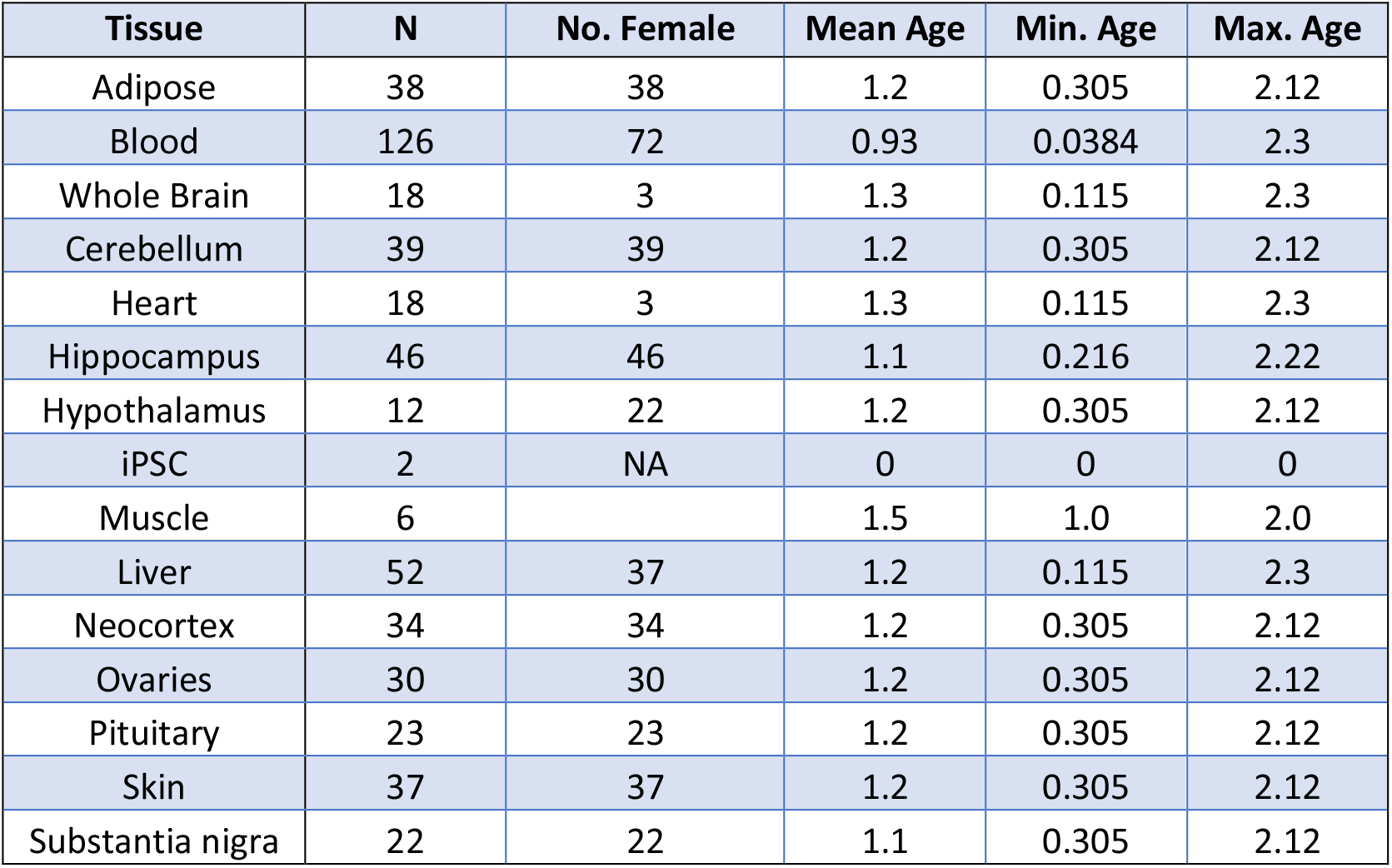
Description of biological materials for training the preliminary epigenetic clocks for rats (N=503). Other supplementary tables provide details on the test data. N=Total number of tissues. Number of females. Age: mean, minimum and maximum in units of years.

**Supplementary Table S2.** Description of the Sprague Dawley rat samples provided by Dr Kavita Singh for training of rat clock.

| Sr. No | Age in Weeks | Age in Months | Number of Animal gender-wise (n=6) | Blood sample | Tissue samples | Total Samples |
| --- | --- | --- | --- | --- | --- | --- |
| 1 | 3 | 0.69 | 3- Male and 3- Female | 6 | - | 6 |
| 2 | 6 | 1.38 | 3- Male and 3- Female | 6 | Heart, Liver, Brain (18 samples) | 24 |
| 3 | 12 | 2.76 | 3- Male and 3- Female | 6 | - | 6 |
| 4 | 28 | 6.44 | 6- Male | 6 | - | 6 |
| 5 | 52 | 12.0 | 6- Male | 6 | Heart, Liver, Brain (18 samples) | 24 |
| 6 | 78 | 18.0 | 3- Male and 3- Female | 6 |  | 6 |
| 7 | 120 | 27.6 | 6- Male | 6 | Heart, Liver, Brain (18 samples) | 24 |
| Total samples (42+54=96) |  |  |  | 42 | 54 | 96 |

**Supplementary Table S3.** Description of the Sprague Dawley rats employed for testing the effects of the E5 plasma fraction treatment. The two set of independent experiments involved different sets of animals. Experiment 1 profiled multiple organs and tissues from male rats. Experiment 2 analyzed blood methylation data from both male and female rats. Experiment 2 was a longitudinal study based on two blood draws: baseline (before any treatment) and 2 weeks post treatment (E5 treatment or saline control).

| Experiment | Group | Age in Weeks | Age in Months | Number of Animal (n=6) | Blood sample | Tissue samples | Total Samples |
| --- | --- | --- | --- | --- | --- | --- | --- |
| 1 | Old Control | 109 | 25.0 | 6 Male | 6+2 replicates | Heart, Liver, Hypothalamus (6 samples each) | 26 |
| 1 | Old Treatment | 109 | 25.0 | 6 Male | 6+2 replicates | Heart, Liver, Hypothalamus (6 samples each) | 26 |
| 1 | Young Control | 30 | 6.9 | 6 Male | 6 | Heart, Liver, Hypothalamus (6 samples each) | 24 |
| **Total samples (22+54=76)** |  |  |  |  | **22** | **54** | **76** |

**Supplementary Table S3.** Description of the Sprague Dawley rats employed for testing the effects of the E5 plasma fraction treatment.
| Experiment | Group | Age in Weeks | Age in Months | Number of Animal | Blood sample | Samples | Total Samples |
| --- | --- | --- | --- | --- | --- | --- | --- |
| 2 | Old Control | 113.4 | 26.0 | 9 (5F, 4M) | 2 per animal | Blood draws separated by 2 weeks | 18 |
| 2 | Old Treated | 113.4 | 26.0 | 8 (6F,2M) | 2 per animal | Blood draws separated by 2 weeks | 16 |
| **Total samples** |  |  |  |  |  |  | **34** |

**Supplementary Table S4:** Mean values of the DNAm age estimates of the six epigenetic clocks in the E5 plasma treatment study. The columns report the figure panels of the corresponding barplot. The tissue that was used to evaluate the clock. The type of clock. The percentage of rejuvenation (last column) was calculated as follows: 100*(1-Old Treated/OldControl). The results for Figure 2 involved the preliminary versions of the six rat clocks. The results for Supplementary Figure 7 involved the final versions of the six rat clocks as detailed in Methods. According to the 6 epigenetic clocks, the plasma fraction treatment rejuvenated liver by 74.6% (ranging from 68.6% to 78.6% depending on the clock), blood by 64.3% (ranging from 52.5 to 74.5%), heart by 46.5%, and hypothalamus by 24.4%. The rejuvenation effects are even more pronounced if we use the final versions of our epigenetic clocks: liver 77.6%, blood 68.2%, heart 56.5%, hypothalamus 29.6%. According to the final version of the epigenetic clocks, the average rejuvenation across four tissues was 67.40%.

| Figure | Tissue | Clock | Old Ctrl | Young Ctrl | Old Treated | Perc. Rejuv. | Ave. Rejuv. By Tissue |
| --- | --- | --- | --- | --- | --- | --- | --- |
| Fig2B | Liver | DNAmAge Pan Tissue | 2.1322 | 0.5260 | 0.5715 | 73.20% |  |
| Fig2F | Liver | DNAmAge Liver | 2.4875 | 0.7220 | 0.7809 | 68.61% |  |
| Fig2J | Liver | DNAmAge HumanRat Pan Tissue | 4.2965 | 1.0316 | 0.9394 | 78.14% |  |
| Fig2N | Liver | DNAmRelativeAge HumanRat Pan Tissue | 0.6752 | 0.2149 | 0.1444 | 78.62% | 74.64% |
| SuppFig12B | Liver | DNAmAge Pan Tissue | 2.1818 | 0.5057 | 0.4997 | 77.10% |  |
| SuppFig12F | Liver | DNAmAge Liver | 2.0457 | 0.5955 | 0.6243 | 69.48% |  |
| SuppFig12J | Liver | DNAmAge HumanRat Pan Tissue | 2.1661 | 0.5847 | 0.4583 | 78.84% |  |
| SuppFig12N | Liver | DNAmRelativeAge HumanRat Pan Tissue | 0.5575 | 0.1562 | 0.0848 | 84.79% | 77.55% |
| Fig2D | Hypothalamus | DNAmAge Pan Tissue | 1.6158 | 0.6597 | 1.2900 | 20.16% |  |
| Fig2H | Hypothalamus | DNAmAge Brain | 2.3387 | 0.9713 | 1.9615 | 16.13% |  |
| Fig2L | Hypothalamus | DNAmAge HumanRat Pan Tissue | 2.1391 | 0.7041 | 1.4740 | 31.09% |  |
| Fig2P | Hypothalamus | DNAmRelativeAge HumanRat Pan Tissue | 0.4861 | 0.1879 | 0.3392 | 30.22% | 24.40% |
| SuppFig12D | Hypothalamus | DNAmAge Pan Tissue | 1.9749 | 0.6954 | 1.4656 | 25.79% |  |
| SuppFig12H | Hypothalamus | DNAmAge Brain | 2.0678 | 0.6363 | 1.5903 | 23.09% |  |
| SuppFig12L | Hypothalamus | DNAmAge HumanRat Pan Tissue | 2.1123 | 0.6277 | 1.3340 | 36.84% |  |
| SuppFig12P | Hypothalamus | DNAmRelativeAge HumanRat Pan Tissue | 0.5256 | 0.1646 | 0.3532 | 32.81% | 29.64% |
| Fig2C | Heart | DNAmAge Pan Tissue | 1.6065 | 0.8285 | 0.8086 | 49.66% |  |
| Fig2G | Heart | DNAmAge Blood | 0.7898 | 0.4014 | 0.4133 | 47.67% |  |
| Fig2K | Heart | DNAmAge HumanRat Pan Tissue | 3.8106 | 1.7115 | 1.9973 | 47.59% |  |
| Fig2O | Heart | DNAmRelativeAge<br>HumanRat Pan<br>Tissue | 0.5783 | 0.2598 | 0.3409 | 41.06% | 46.49% |
| SuppFig12C | Heart | DNAmAge Pan<br>Tissue | 1.9428 | 0.6844 | 0.8074 | 58.44% |  |
| SuppFig12G | Heart | DNAmAge Blood | 0.7704 | 0.4283 | 0.4367 | 43.32% |  |
| SuppFig12K | Heart | DNAmAge<br>HumanRat Pan<br>Tissue | 2.0572 | 0.6036 | 0.7483 | 63.62% |  |
| SuppFig12O | Heart | DNAmRelativeAge<br>HumanRat Pan<br>Tissue | 0.5471 | 0.1613 | 0.2157 | 60.57% | 56.49% |
| Fig2A | Blood | DNAmAge Pan<br>Tissue | 1.7767 | 0.7497 | 0.8435 | 52.53% |  |
| Fig2E | Blood | DNAmAge Blood | 2.1596 | 0.6614 | 0.7867 | 63.57% |  |
| Fig2I | Blood | DNAmAge<br>HumanRat Pan<br>Tissue | 1.7369 | 0.6152 | 0.4429 | 74.50% |  |
| Fig2M | Blood | DNAmRelativeAge<br>HumanRat Pan<br>Tissue | 0.4724 | 0.1777 | 0.1585 | 66.46% | 64.26% |
| SuppFig12A | Blood | DNAmAge Pan<br>Tissue | 1.9947 | 0.6744 | 0.7830 | 60.75% |  |
| SuppFig12E | Blood | DNAmAge Blood | 1.9523 | 0.6094 | 0.7385 | 62.18% |  |
| SuppFig12I | Blood | DNAmAge<br>HumanRat Pan<br>Tissue | 2.0081 | 0.5998 | 0.4395 | 78.11% |  |
| SuppFig12M | Blood | DNAmRelativeAge<br>HumanRat Pan<br>Tissue | 0.5394 | 0.1630 | 0.1533 | 71.58% | 68.15% |

**Supplementary Table S5:** Measurement (with standard deviations) of grip strength of indicated groups of 6 rats each. These average values formed the graphs in Supplementary Figure 9B.

| Groups | Grip Strength in N |  |  |
| --- | --- | --- | --- |
|  | Old control | Treatment | Adult Control |
| **0 Day** | 6.10±0.32 | 6.31±0.44 | 10.25±1.01 |
| **4 Day** | 6.24±0.44 | 8.25±0.94 | 10.84±1.12 |
| **8 Day** | 6.01±0.46 | 11.38±0.83 | 10.55±0.99 |
| **15 Day** | 6.00±0.89 | 11.55±0.88 | 11.35±1.15 |
| **30 Day** | 5.78±0.75 | 11.74±0.76 | 12.01±1.23 |

**Supplementary Table S6:**
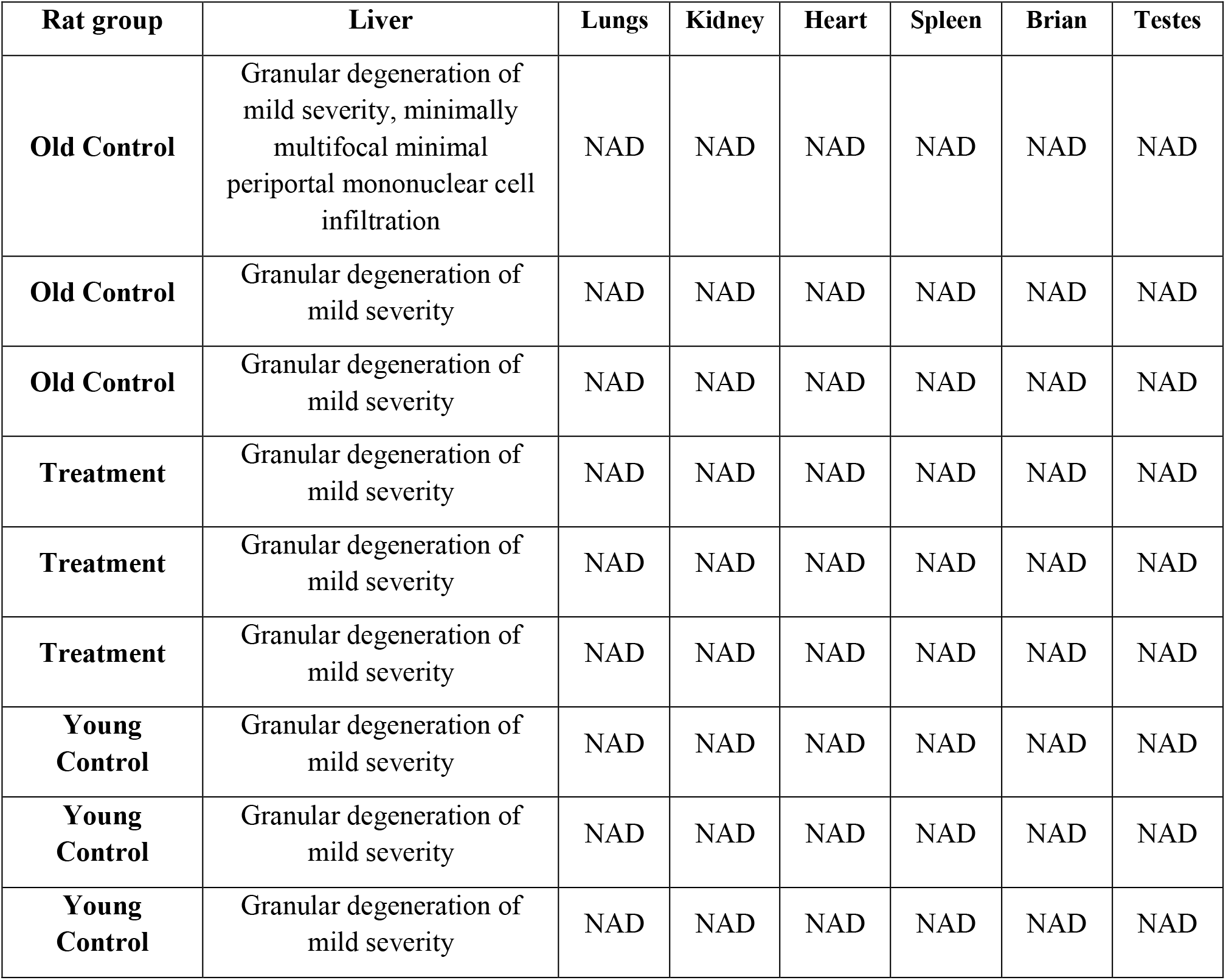
Results of histopathological analyses of rat tissues. Lesions suggestive of any toxicity were not noted. NAD= No Abnormalities Detected. Representative images of the tissues are shown in Supplementary Figure 9.

| Rat group | Liver | Lungs | Kidney | Heart | Spleen | Brian | Testes |
| --- | --- | --- | --- | --- | --- | --- | --- |
| Old Control | Granular degeneration of mild severity, minimally multifocal minimal periportal mononuclear cell infiltration | NAD | NAD | NAD | NAD | NAD | NAD |
| Old Control | Granular degeneration of mild severity | NAD | NAD | NAD | NAD | NAD | NAD |
| Old Control | Granular degeneration of mild severity | NAD | NAD | NAD | NAD | NAD | NAD |
| Treatment | Granular degeneration of mild severity | NAD | NAD | NAD | NAD | NAD | NAD |
| Treatment | Granular degeneration of mild severity | NAD | NAD | NAD | NAD | NAD | NAD |
| Treatment | Granular degeneration of mild severity | NAD | NAD | NAD | NAD | NAD | NAD |
| Young Control | Granular degeneration of mild severity | NAD | NAD | NAD | NAD | NAD | NAD |
| Young Control | Granular degeneration of mild severity | NAD | NAD | NAD | NAD | NAD | NAD |
| Young Control | Granular degeneration of mild severity | NAD | NAD | NAD | NAD | NAD | NAD |

**Supplementary Table S7:**
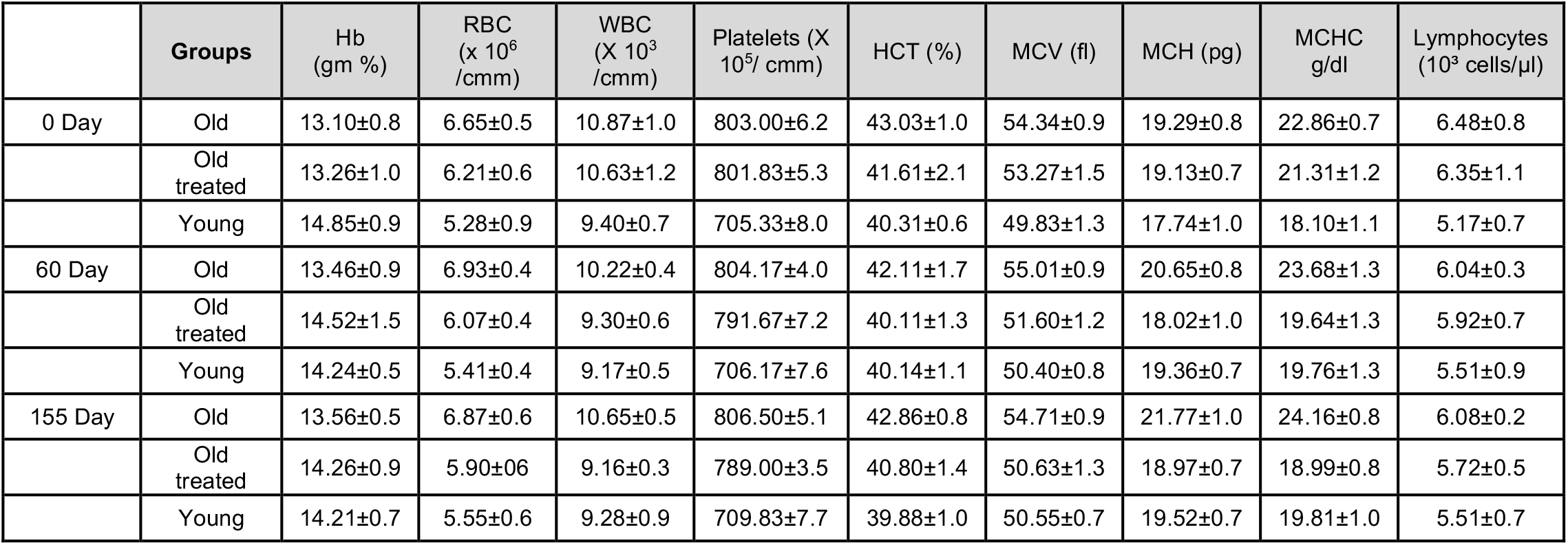
Blood indices measurements (with standard deviation) at indicated time points post-treatment. Each measurement was taken from 6 rats per group. These average values formed the graphs in Supplementary Figure 12.

| | Groups | Hb<br>(gm %) | RBC<br>(x 10<sup>6</sup><br>/cmm) | WBC<br>(X 10<sup>3</sup><br>/cmm) | Platelets (X<br>10<sup>5</sup>/ cmm) | HCT (%) | MCV (fl) | MCH (pg) | MCHC<br>g/dl | Lymphocytes<br>(10<sup>3</sup> cells/μl) |
| --- | --- | --- | --- | --- | --- | --- | --- | --- | --- | --- |
| 0 Day | Old | 13.10±0.8 | 6.65±0.5 | 10.87±1.0 | 803.00±6.2 | 43.03±1.0 | 54.34±0.9 | 19.29±0.8 | 22.86±0.7 | 6.48±0.8 |
| | Old<br>treated | 13.26±1.0 | 6.21±0.6 | 10.63±1.2 | 801.83±5.3 | 41.61±2.1 | 53.27±1.5 | 19.13±0.7 | 21.31±1.2 | 6.35±1.1 |
| | Young | 14.85±0.9 | 5.28±0.9 | 9.40±0.7 | 705.33±8.0 | 40.31±0.6 | 49.83±1.3 | 17.74±1.0 | 18.10±1.1 | 5.17±0.7 |
| 60 Day | Old | 13.46±0.9 | 6.93±0.4 | 10.22±0.4 | 804.17±4.0 | 42.11±1.7 | 55.01±0.9 | 20.65±0.8 | 23.68±1.3 | 6.04±0.3 |
| | Old<br>treated | 14.52±1.5 | 6.07±0.4 | 9.30±0.6 | 791.67±7.2 | 40.11±1.3 | 51.60±1.2 | 18.02±1.0 | 19.64±1.3 | 5.92±0.7 |
| | Young | 14.24±0.5 | 5.41±0.4 | 9.17±0.5 | 706.17±7.6 | 40.14±1.1 | 50.40±0.8 | 19.36±0.7 | 19.76±1.3 | 5.51±0.9 |
| 155 Day | Old | 13.56±0.5 | 6.87±0.6 | 10.65±0.5 | 806.50±5.1 | 42.86±0.8 | 54.71±0.9 | 21.77±1.0 | 24.16±0.8 | 6.08±0.2 |
| | Old<br>treated | 14.26±0.9 | 5.90±0.6 | 9.16±0.3 | 789.00±3.5 | 40.80±1.4 | 50.63±1.3 | 18.97±0.7 | 18.99±0.8 | 5.72±0.5 |
| | Young | 14.21±0.7 | 5.55±0.6 | 9.28±0.9 | 709.83±7.7 | 39.88±1.0 | 50.55±0.7 | 19.52±0.7 | 19.81±1.0 | 5.51±0.7 |

**Supplementary Table S8:**
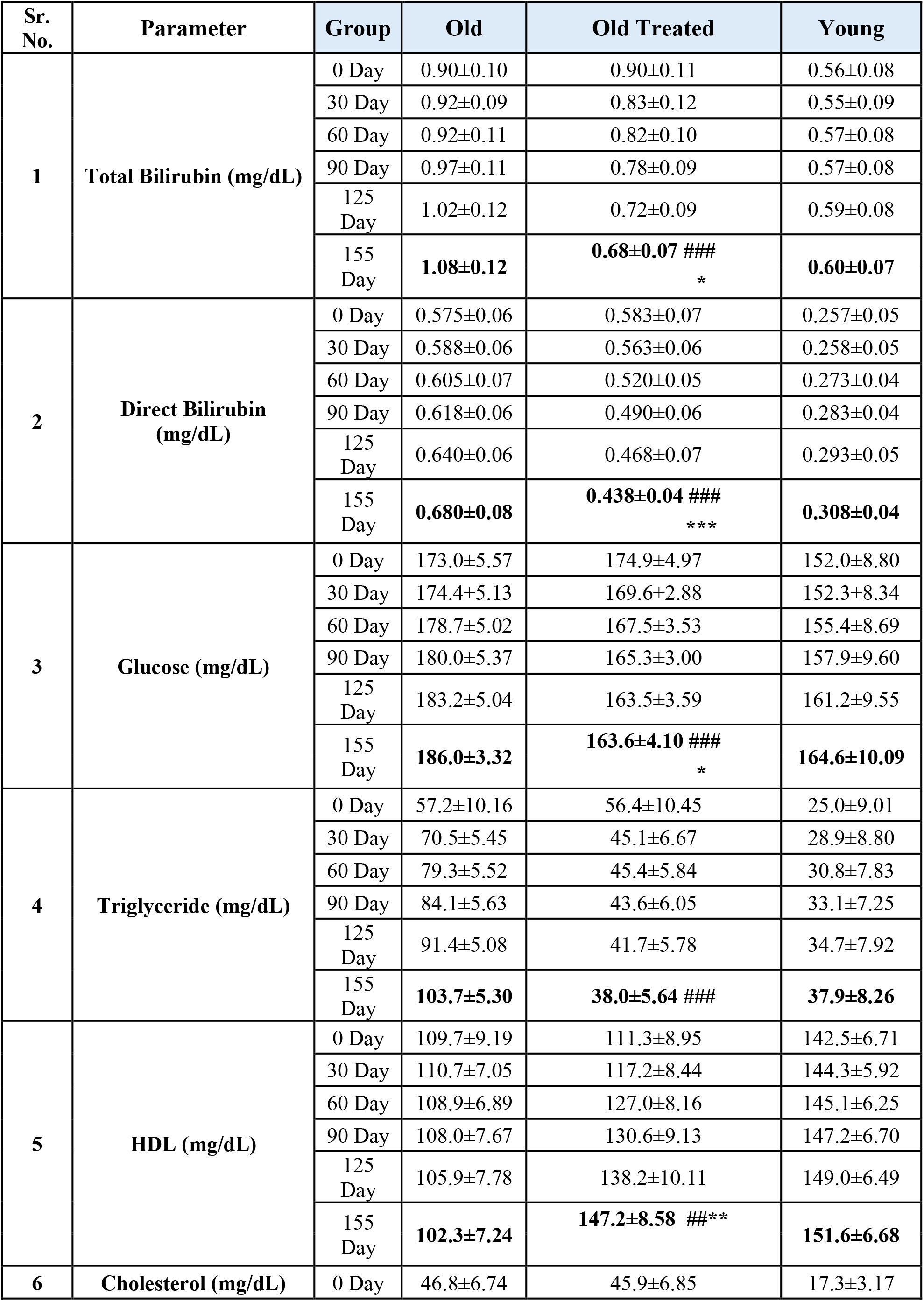
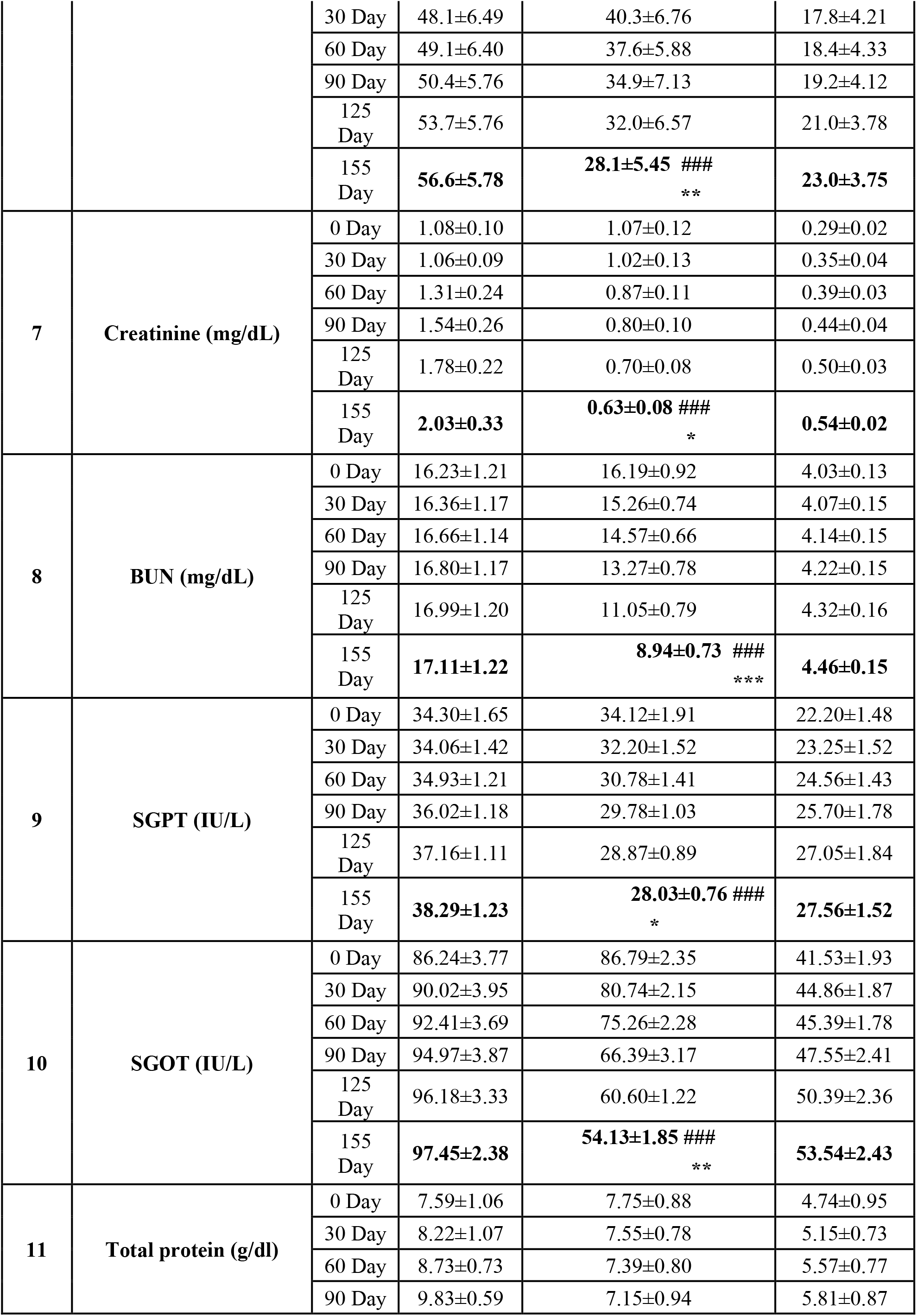
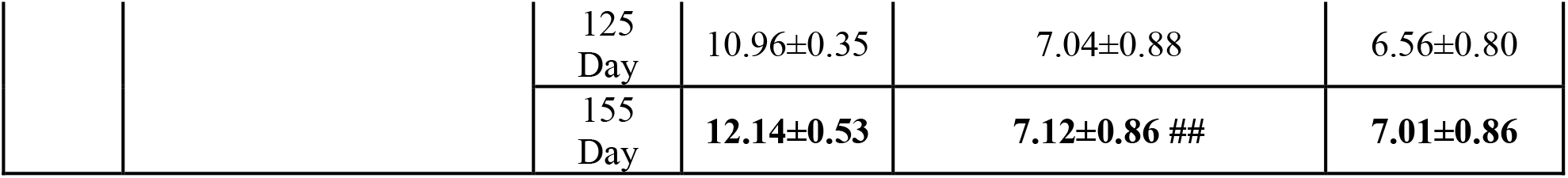
Detailed vital organ biomarker measurements of rats at stated time points post-plasma fraction treatment. The measurements (with standard deviation) were from 6 rats per group. These average values formed the graphs in Figure 3.

**Supplementary Table S9.**
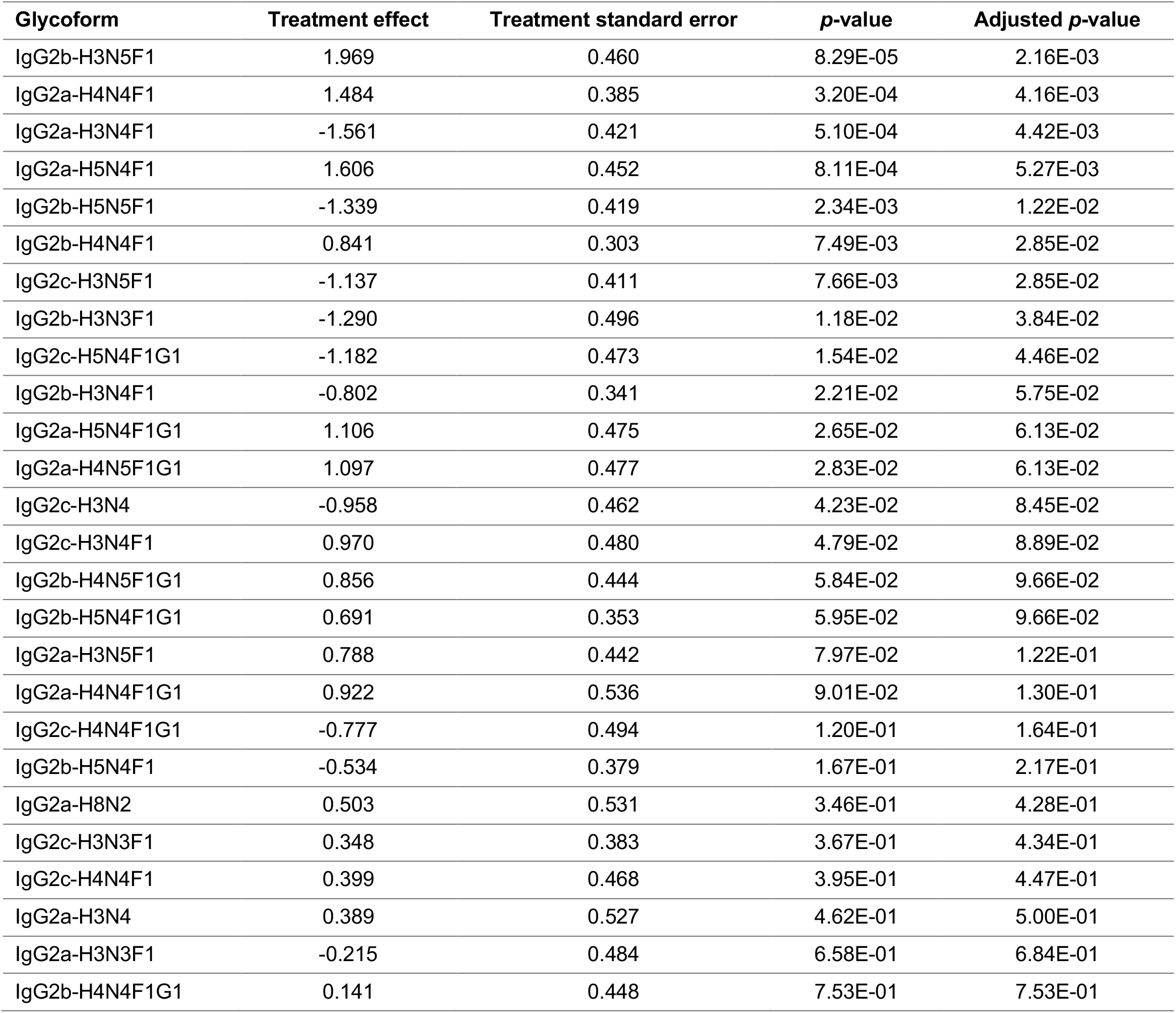
Glycan age analysis in rat blood. Plasma fraction E5 treatment effect on rat IgG2a, 2b, and 2c N-glycoform composition. Glycoproteomic data was analyzed using a linear mixed-effects model and p-values < 0.05 were considered statistically significant. The rows correspond to different rat IgG tryptic peptides as detailed in the following panel.

## Technical Details and software for the rat clocks

The epigenetic clock software for rats can be applied to data generated on the mammalian array platform ^1^. New data for epigenetic clock studies can be generated with the mammalian methylation array (HorvathMammalMethylChip40), which is distributed by the Epigenetic Clock Development Foundation: https://clockfoundation.org/

The coefficient values and CpGs underlying the clocks can be found in a Supplementary Table.

### Statistical methods used for building the clocks

The epigenetic clocks were used by employing a single elastic net regression model analysis (R function glmnet). We use used Leave-one-out analysis (LOO) using a single lambda value. We chose the following parameters for the glmnet R function (Alpha: 0.5, CV Fold: 10, Lambda choice for Clock: 1 standard error above minimum CV-MSE).

We report 2 sets of clocks (preliminary versions and final versions). We recommend to use the final versions. We only report the preliminary versions so that readers can recreate Figure 1 and Figure 2 in our article.

### Clocks

1) The preliminary and final version of the pan tissue clock for rats is based on 195 CpGs and 193 CpGs which are specified in columns “Coef.PrelimRatPanTissue” and “Coef.FinalRatPanTissue”, respectively. No age transformation was carried out.

2) The preliminary and final version of the blood tissue clock for rats is based on 48 CpGs and 51 CpGs which are specified in columns “Coef.PrelimRatBloodTissue” and “Coef.FinalRatBloodTissue”, respectively. No age transformation was carried out.

3) The preliminary and final version of the Brain tissue clock for rats is based on 99 CpGs and 108 CpGs which are specified in columns “Coef.PrelimRatBrainTissue” and “Coef.FinalRatBrainTissue”, respectively. No age transformation was carried out.

4) The preliminary and final version of the Liver tissue clock for rats is based on 62 CpGs and 46 CpGs which are specified in columns “Coef.PrelimRatLiverTissue” and “Coef.FinalRatLiverTissue”, respectively. No age transformation was carried out.

5) Preliminary and final versions of the human rat pan tissue clock are specified in Coef.PrelimHumanRatPanTissueLogLinearAge (700 CpGs) and Coef.FinalHumanRatPanTissueLogLinearAge (989 CpGs). Note that a log linear transformation was applied.

6) Preliminary and final versions of the human rat blood tissue clock are specified in Coef.PrelimHumanRatBloodTissueLogLinearAge (97 CpGs) and Coef.FinalHumanRatBloodTissueLogLinearAge (72 CpGs). Note that a log linear transformation was applied.

7) Preliminary and final versions of the human rat pan tissue clock for RELATIVE AGE are specified in Coef.PrelimHumanRatRelativeAgePan (870 CpGs) and Coef.FinalHumanRatRelativeAgePan (738 CpGs). Note that we define RelativeAge=Age/maxLifespan where maxLifespan is 122.5 for humans and 3.8 years for rats, respectively. Age is in units of years.

8) Preliminary and final versions of the human rat blood tissue clock for RELATIVE AGE are specified in Coef.PrelimHumanRatRelativeAgeBlood (138 CpGs) and Coef.FinalHumanRatRelativeAgeBlood (145 CpGs).

The DNAm Age estimate is estimated as follows. Form a weighted linear combination of the CpGs whose details can be found in the Supplementary File, SupplementaryData.RatClockCoef.csv.

This will result in a number referred to as LinearCombination. Some of the clocks involve a log linear transformation whose inverse needs to be applied.

The Supplementary file reports the probe identifier (cg number) used in the custom Infinium array (HorvathMammalMethylChip40). The weights used in this linear combination are specified in the respective column entitled “Coef.”.

The formula assumes that the DNA methylation data measure “beta” values, but the formula could be adapted to other ways of generating DNA methylation data.

### General description of age transformation

The human-rat clocks for chronological age used log linear transformations that are similar to those employed for the HUMAN pan tissue (Horvath 2013, Genome Biology).

An elastic net regression model (implemented in the glmnet R function) was used to regress a transformed version of age on the beta values in the training data. The glmnet function requires the user to specify two parameters (alpha and beta). Since we used an elastic net predictor, alpha was set to 0.5. But the lambda value of was chosen by applying a 10 fold cross validation to the training data (via the R function cv.glmnet).

The elastic net regression results in a linear regression model whose coefficients b_0_, b_1_, . . ., relate to transformed age as follows

*F*(chronological age)=*b*_0_*+b*_1_*CpG*_1_*+ . . . +b*_p_*CpG*_p_+error

Note that the intercept term is denoted by b_0_. The coefficient values can be found in the Supplementary file. Based on the coefficient values from the regression model, DNAmAge is estimated as follows

*DNAm*Age=𝐹^−1^(*b*_0_*+b*_1_*CpG*_1_*+ . . . +b*_p_*CpG*_p_)

, where 𝐹^−1^(𝑦) denotes the mathematical inverse of the function F(.). Thus, the regression model can be used to predict to transformed age value by simply plugging the beta values of the selected CpGs into the formula.

### Defining Properties of the log linear transformation

As indicated by its name, the “log-linear” function, has a logarithmic dependence on age before the average age of sexual maturity (of the species) and a linear dependence after Age at Sexual Maturity (of the species). For the human-rat clocks we used the following averages at sexual maturity (in units of years): 13.5 years for humans and 0.219178082 years for rats (Supplementary Table).

We used a piecewise transformation, parameterized by Age of Sexual Maturity (𝐴). The transformation is F(x), given by

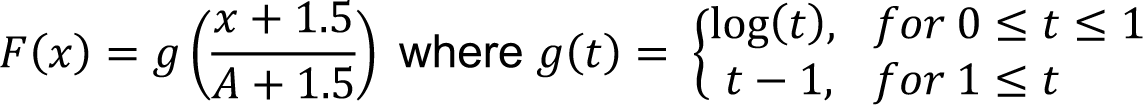

Explicitly, F(x) is given by

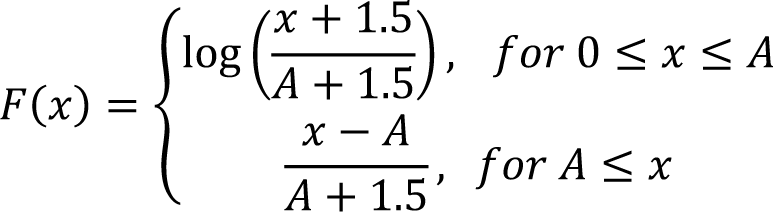

In order to use this transformation to predict Age on *new samples*, one needs to use the *inverse* transformation, F^-1^(y), given by

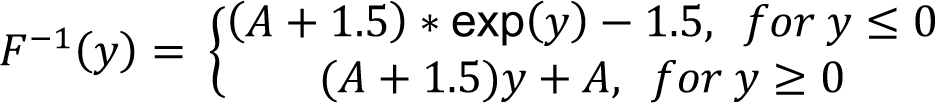

For predicting age, apply the inverse transformation to coefficient-weighted sum. That is,

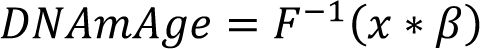

where 𝛽 is the vector of coefficients and 𝑥 is the vector of methylation values, with an intercept term.

### R Implementation of the log linear transformation

\### Applies the log linear transformation to the input vector x, i.e. to Age

F= Vectorize(function(x, maturity, …) {

if (is.na(x) | is.na(maturity)) {return(NA)} k <-1.5

y <-0

if (x < maturity) {y = log((x+k)/(maturity+k))} else {y = (x-maturity)/(maturity+k)}

return(y)

})

\### Inverse log linear transformation

F.inverse= Vectorize(function(y, maturity, …) { if (is.na(y) | is.na(maturity)) {return(NA)}

k <-1.5

x <-0

if (y < 0) {x = (maturity+k)*exp(y)-k} else {x = (maturity+k)*y+maturity} return(x)

})

### The DNAm Age estimate is estimated in two steps

First, one forms a weighted linear combination of the CpGs whose details can be found in Table

The table reports the probe identifier (cg number) used in the custom Infinium array (HorvathMammalMethylChip40). The weights used in this linear combination are specified in the respective column entitled “Coef.”.

The formula assumes that the DNA methylation data measure “beta” values but the formula could be adapted to other ways of generating DNA methylation data.

Pseudo R code

\# R function for multivariate regression model multivariatePredictorCoef=function(dat0, datCOEF,imputeValues=FALSE)

{

datout=data.frame(matrix(NA,nrow=dim(dat0)[[2]]- 1,ncol=dim(datCOEF)[[2]]-1)) match1=match(datCOEF[-1,1],dat0[,1])

if (sum(!is.na(match1))==0) stop(“Input error. The first column of dat0 does not contain CpG identifiers (cg numbers).”) dat1=dat0[match1,]

row.names1=as.character(dat1[,1]) dat1=dat1[,-1]

if (imputeValues){dat1=impute.knn(data=as.matrix(dat1),k = 10)[[1]]}

for (i in 1:dim(dat1)[[2]]){ for (j in

2:dim(as.matrix(datCOEF))[[2]]){

datout[i,j-1]=sum(dat1[,i]* datCOEF[-1,j],na.rm=TRUE)+ datCOEF[1,j]}}

colnames(datout)=colnames(datCOEF)[-1] rownames(datout)=colnames(dat0)[-1] datout=data.frame(SampleID= colnames(dat0)[-1],datout) datout

} # end of function

\# read in supplementary table datCoef=read.csv(“TableS.csv”)

The first columns should read as follows names(datCoef)

1) var

2) Coef.PrelimRatPanTissue

3) Coef.PrelimRatBloodTissue

4) Coef.PrelimRatBrainTissue

5) Coef.PrelimRatLiverTissue

6) Coef.PrelimHumanRatPanTissueLogLinearAge

7) Coef.PrelimHumanRatBloodTissueLogLinearAge

8) Coef.PrelimHumanRatRelativeAgePan

9) Coef.PrelimHumanRatRelativeAgeBlood

10) Coef.FinalRatPanTissue

11) Coef.FinalRatBloodTissue

12) Coef.FinalRatBrainTissue

13) Coef.FinalRatLiverTissue

14) Coef.FinalHumanRatPanTissueLogLinearAge

15) Coef.FinalHumanRatBloodTissueLogLinearAge

16) Coef.FinalHumanRatRelativeAgePan

17) Coef.FinalHumanRatRelativeAgeBlood ETC ETC

\# Restrict attention to the FINAL rat clocks columns datCoef=datCoef[,c(1,10:17)]

\# assume the first column of dat0 contains the CpG identifiers match1=match(datCoef[-1,1],dat0[,1])

missingProbes= as.character(datCoef[-1,1])[is.na(match1)]

dat1=dat0[match1,]

\# data frame with predicted values. datPredictions=multivariatePredictorCoef(dat1,datCOEF=datCoef,impute Values=FALSE)

#let’s relabel the columns by replacing “Coef” with “DNAm” since the columns contain estimates of age or relative age instead of coefficient values

colnames(datPredictions)=gsub(pattern=“Coef”, replacement=“DNAm”, x=colnames(datPredictions))

\# We need to transform the human rat clock for chronological age using the inverse of the log linear #transformation.

For rats, the age at sexual maturity has to be set to 0.21917808 years. datPredictions$DNAm.FinalHumanRatPanTissueLogLinearAge=

F.inverse(datPredictions$DNAm.FinalHumanRatPanTissueLogLinearAge, maturity=0.21917808) datPredictions$DNAm.FinalHumanRatBloodTissueLogLinearAge= F.inverse(datPredictions$DNAm.FinalHumanRatBloodTissueLogLinearAge, maturity=0.21917808)

#The data frame “datPredictions” contains the age estimates in units of years and relative age estimates.

